# A Bayesian approach to accurate and robust signature detection on LINCS L1000 data

**DOI:** 10.1101/769620

**Authors:** Yue Qiu, Tianhuan Lu, Hansaim Lim, Lei Xie

## Abstract

LINCS L1000 dataset produced by L1000 assay contains numerous cellular expression data induced by large sets of perturbagens. Although it provides invaluable resources for drug discovery as well as understanding of disease mechanisms, severe noise in the dataset makes the detection of reliable gene expression signals difficult. Existing methods for the peak deconvolution, either *k*-means based or Gaussian mixture model, cannot reliably recover the accurate expression level of genes in many cases, thereby limiting their robust applications in biomedical studies. Here, we have developed a novel Bayes’ theory based deconvolution algorithm that gives unbiased likelihood estimations for peak positions and characterizes the peak with a probability based *z*-scores. Based on above algorithm, a pipeline is built to process raw data from L1000 assay into signatures that represent the features of perturbagen. The performance of the proposed new pipeline is rigorously evaluated using the similarity between bio-replicates and between drugs with shared targets. The results show that the new signature derived from the proposed algorithm gives a substantially more reliable and informative representation for perturbagens than existing methods. Thus, our new Bayesian-based peak deconvolution and *z*-score calculation method may significantly boost the performance of invaluable L1000 data in the down-stream applications such as drug repurposing, disease modeling, and gene function prediction.

## 1 Introduction

The Library of Integrated Network-based Cellular Signatures (LINCS) created a resource of cellular state changes under treatment of perturbagens, including chemical compounds, RNAis, and CRISPRs.^1^ L1000 assay is used in LINCS as a large-scale gene expression profiling assay, which provides low cost, high throughput gene expression profiling on perturbagen treated cells.^2^ The application of L1000 assay allows the profiling on more than one million samples treated with more than 50,000 different perturbagens across 98 cell lines. These profiles are processed into molecular signatures to represent cellular effects of certain perturbagen treatment.^3^ This comprehensive profiling provided by L1000 is widely used in drug discovery and repurposing, largely facilitating large scale pharmacology analysis.^4, 5^

L1000 assay measures the expression of 978 landmark genes with The Luminex FlexMap 3D platform, which can identify 500 different bead color as tags for different genes. To measure all landmark genes within one scan, L1000 separately coupled two different gene barcodes to aliquots of the same bead color and mixed them with a ratio of 2:1. In consequence, two peaks in the distribution of fluorescent intensity (FI) are expected, and a deconvolution step must be involved to access the expression level of a certain gene. LINCS adopted the *k*-means clustering algorithm to separate all reads of the same beads color into two distinct components, and the median fluorescent intensity values are assigned to each gene.

Although the *k*-means clustering gives a good estimation on most of the data, cases with unexpected ratio between two peaks, classified into more than two categories, or large overlap between peaks can not be well solved.^6^ This problem limits the quality of *z*-score profiles, and adds to the difficulty of utilizing the massive data provided by L1000 assay.

After L1000 data was released, efforts have been made to improve the peak deconvolution process. Liu et al. developed a method based on Gaussian mixture model (GMM) to improve the accuracy of peak deconvolution.^7^ They assume that each gene’s fluorescent intensities follow a Gaussian distribution and thus each sample with two genes will subject to a bimodal Gaussian distribution. GMM can avoid over-clustering problem in *k*-means but raises a new problem that the *z*-scores are highly sensitive to frequent isolated reads and become unreliable. To solve this problem, Li et al. developed an aggregate Gaussian mixture model (AGMM) and associate software (l1kdeconv).^8^ They added an outlier identification step before deconvoluting peaks with GMM to make the algorithm more robust. However, there are still cases where the peaks cannot be well identified. Thus, it is necessary to develop a new algorithm that improves the data process of L1000 assay.

In this study, we describe a novel peak deconvolution algorithm based on Bayes’ theorem and a probability based *z*-score inference method. We model each measurement as a random sample from the population with two different components mixed by 2:1 ratio. The likelihoods for all different fluorescent intensity values are calculated by Bayes’ theorem. Then *z*-scores are inferred by the probabilities for the genes to have differential expression. The gene expression profiles deconvoluted from our Bayesian method achieve higher similarity between bio-replicates and drugs with shared targets than those generated from the existing methods. This suggests that the molecular signatures from our method are of better consistency and less noisy, which will be helpful for further pharmacology analysis.

We develop a pipeline to process raw data from L1000 assay into *z*-scores. The code and the precomputed data for LINCS L1000 Phase II (GSE 70138) are available at https://github.com/njpipeorgan/L1000-bayesian.

## 2 Results

### 2.1 Work flow

Based on a novel Bayesian analysis-based peak deconvolution and a new *z*-score inference method, we develop a pipeline to generate signature from raw data measured from L1000 assay. The pipeline takes raw fluorescent intensity data from LINCS L1000 datasets as input and gives a combined *z*-score profile for each experiment as its signature. As shown in Fig. 1, our pipeline is composed of 5 steps as follows:

**Figure 1:**
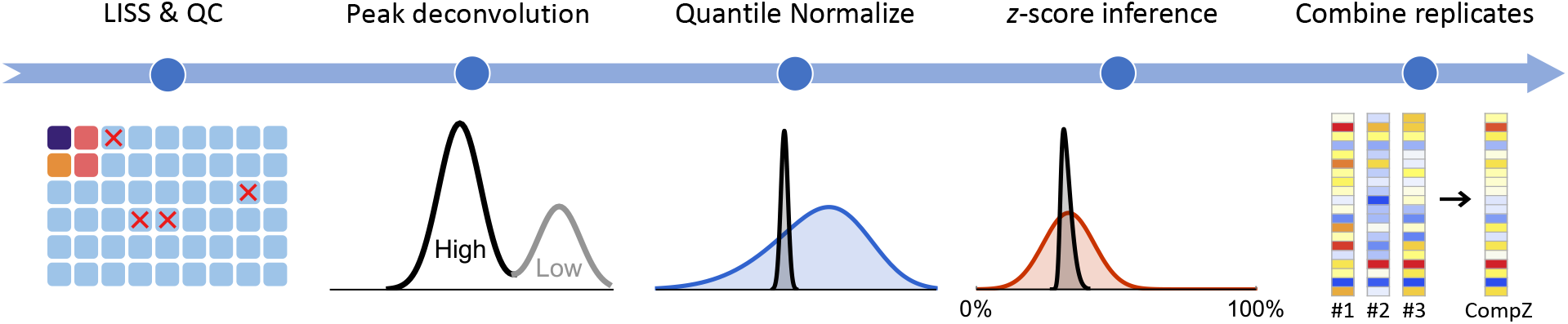
Illustration of pipeline for robust L1000 perturbagen signature detection.

1. LISS and quality control. In this step, we perform a two step linear scaling to calibrate the fluorescent intensities. Also, we identify the low-quality samples with goodness of fit *χ*^2^ > 4.0 or the slope *a* > 3.0.
2. Peak deconvolution. For beads coupled with two different transcript barcodes, a deconvolution step is involved to infer the peak position for each gene. Two probability distributions will be given to the transcripts as the estimations of their expression levels.
3. Quantile normalization. The shape of expression profile is standardized across all samples on the same plate so that different samples on the same plate are comparable to each other.
4. Probablistic *z*-score inference. *z*-scores are inferred by comparing the probability distribution for each gene with its background distribution. They represent relative gene expressions.
5. Combining replicates. *z*-scores profiles from bio-replicates are combined into one signature by weighted average.

### 2.2 Peak deconvolution

We assess our peak deconvolution method by check the consistency between the peak locations from our Bayesian approach, L1000 level 2 data, and AGMM. For our method, since the peak deconvolution process gives probability distributions instead of precise numbers for peak locations, we determine the peak locations here by maximizing their marginal probability distributions (see Online Methods), and it is refer to as Bayesian (MLE) hereafter.

When comparing the locations of a peak by two methods, we consider them to be different when the discrepancy in log2 expression value is larger than 0.2, which is based on the fact that the peaks have a typical scale parameter *σ* of ≳ 0.2. Also, all the methods are considered giving the same location for a peak when all three values are within ±0.2 range relative to the middle one. We compare all 978 peak locations in a sample well for three methods, and the results are shown in Table 1.

**Table 1:** The consistency in peak deconvolution between our Bayesian method, L1000, and AGMM. Peak locations for 978 landmark genes are determined in well REP.A028_MCF7_24H_X2_B25_D11 by each method. The last three cases show those where two of the methods give the same peak location while the third one gives a different location.

| Case | Description | Number of genes |
| --- | --- | --- |
| 1 | All are the same | 773 |
| 2 | All are different | 22 |
| 3 | Only L1000 is different | 86 |
| 4 | Only AGMM is different | 79 |
| 5 | Only Bayesian (MLE) is different | 18 |

In most of the cases, all three methods agrees with each other, which shows an overall consistency among all methods. Although all three methods may give a prediction that different from the othre two, it happens less frequently to Bayesian (MLE) method (18 verses 86 and 79 in Table 1). To further investigate the origin of those disagreements in Table 1, we show one typical example for each case in Fig. 2. A full list of all peak locations by the three methods can be found in supplementary materials. Here, we give a brief analysis for each case:

**Figure 2:**
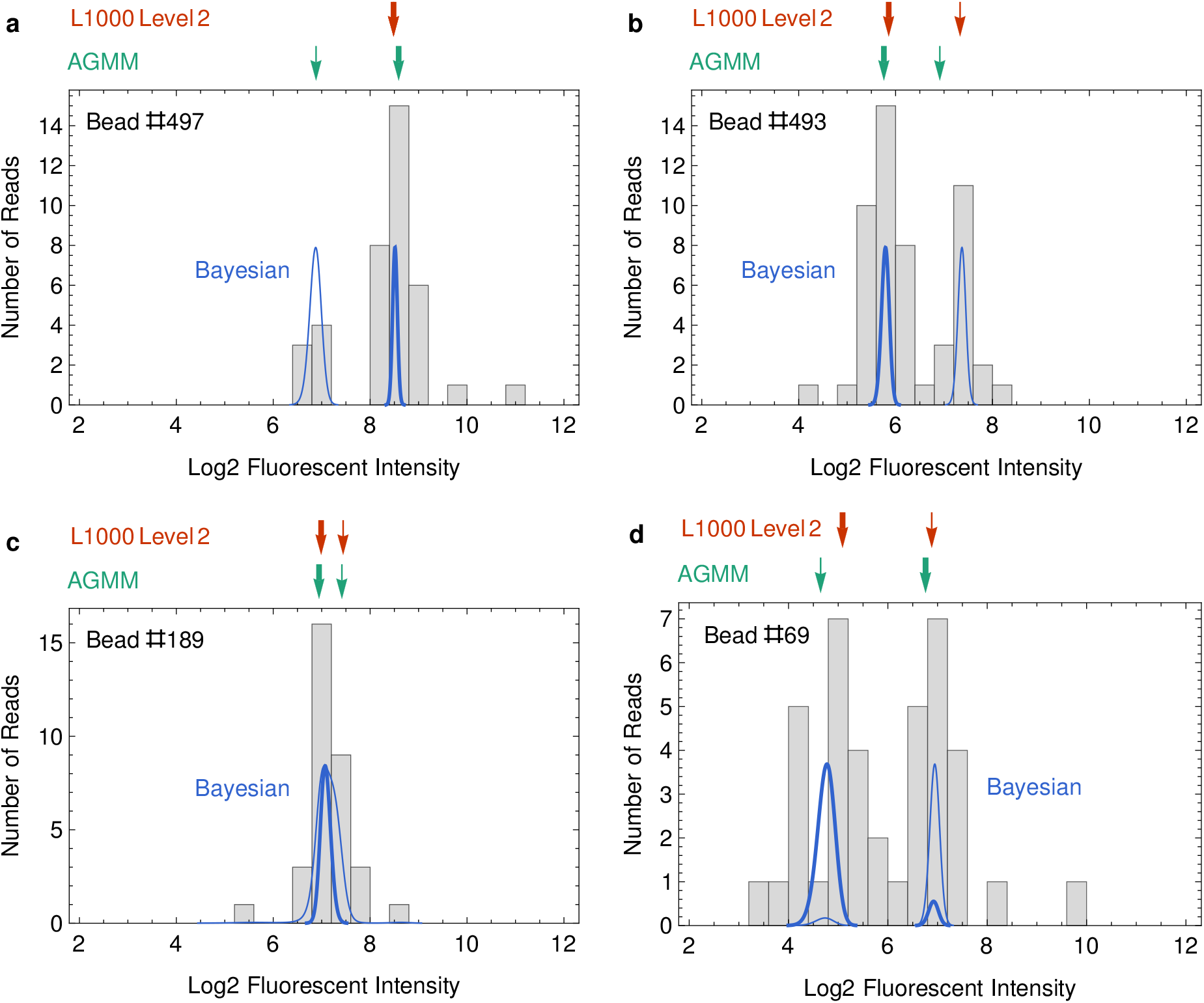
Typical examples where discrepancies happens between MLE, L1000 Level 2 data, and AGMM. For each panel, the results from L1000 peak deconvolution and AGMM are shown in red and green arrows, where the thick and thin arrows indicate the peaks with high (2/3) and low (1/3) abundances respectively. The results from our method are shown as probability distributions for high and low peaks in thick and thin blue curves respectively. The examples are from well REP.A028_MCF7_24H_X2_B25_D11.

- Fig. 2(**a**) is an example for Case 3 in Table 1. *k*-means clustering sometimes fail to give two clusters of reads. In those scenarios, L1000 sets the expression level of both genes to be the median of all reads, which yields a different result from Bayesian (MLE) and AGMM.
- Fig. 2(**b**) is an example for Case 4 in Table 1. AGMM sometimes fails to identify the peaks that are not well separated. In the case of Fig. 2(**b**), AGMM gives a scale parameter *σ* of 0.55, which overestimates the actual peak widths, and it makes AGMM to separate the peaks at a wrong location.
- Fig. 2(**c**) is an example for Case 5 in Table 1. When the two peaks are mostly overlapped, Bayesian (MLE) tends to predict the peaks at roughly the same location, while both L1000 and AGMM tends to give two different peak location. In this case, L1000 and AGMM agree with each other and Bayesian (MLE) gives a different result.
- Fig. 2(**d**) is an example for Case 2 in Table 1. All three methods gives different predictions for the high abundance peak in this case. For L1000, reads with log2 FI smaller than 4 are discarded so the peak location is biased to have a higher expression. For AGMM, the separation of the two peaks is not correct, and the locations of the two peaks are flipped.

From these cases, we find that except when the information is not enough to decide the peak locations well, the Bayesian method always gives reasonable predictions but L1000 level 2 and AGMM sometimes make mistakes. It explains why the Bayesian (MLE) data have a lower chance of being different from two other methods simultaneously. We also find that the rate of disagreement between our method and L1000 is around 10%, which should be considered crucial. Given that the genes regulated under each treatment is very limited, the 10% noise signal will largely affect the quality of the dataset.

### 2.3 Similarity between replicates

After locating peaks from raw FI values, we quantile normalize the data and infer their *z*-scores. The *z*-scores can be interpreted as relative gene expression levels, where significantly up/down-regulated genes will get a large positive/negative *z*-scores. The *z*-scores also serves as the final feature of expression for each sample.

L1000 dataset typically include three bio-replicates for each experiment. Since they are expected to be similar to each other in gene expression, a validation of this statistical property indicates good quality of the data and the processing pipeline. L1000 performs multiple experiments with different doses for each perturbagen. In order to maximize the difference between the perturbagens, we only take those experiments with the maximum doses on each plate in measuring the performance.

In this study, we adopt Gene Set Enrichment Analysis (GSEA) to compare gene expression profiles, which is a non-parametric statistical method widely used in analyzing gene expression data.^11^ GSEA yields a score for the enrichment of a gene set (called a query hereafter) in an expression profile. Here we directly take the *z*-scores as the expression profiles and build the queries by taking an equal number of genes that are most up/down regulated in terms of their *z*-scores. The higher enrichment score a query get, the more similar the query and expression profile are.

Besides L1000 level 4 data, we also use the *z*-scores data by Bayesian (MLE). Where peak position from MLE are obtained in the same way as described in Sec. 2.2, followed by quantile normalization and *z*-score inference follows L1000 methods in the exact ways. As shown in Table 1, MLE can summerize the advantage of L1000 and AGMM, so we use it as an optimized determinstic methods and the control for probablistic *z*-score inference (see Online Methods for details).

We test the performance of the methods as follows. For each sample, we obtain its background distribution of GSEA scores by querying the sample against all other samples in the same cell line. Then we query the sample against one of its bio-replicates. For each false positive rate (FPR), i.e. a certain ratio of the highest background scores being picked, a true positive rate can be calculated as the probability that its bio-replicate has a GSEA score higher than the threshold. Fig. 3(**a**) shows the performance of all three methods with different query sizes. Among all query size we tested, our Bayesian method has higher TPR at the same FPR and the performance difference is the most significant when the query size is small.

**Figure 3:**
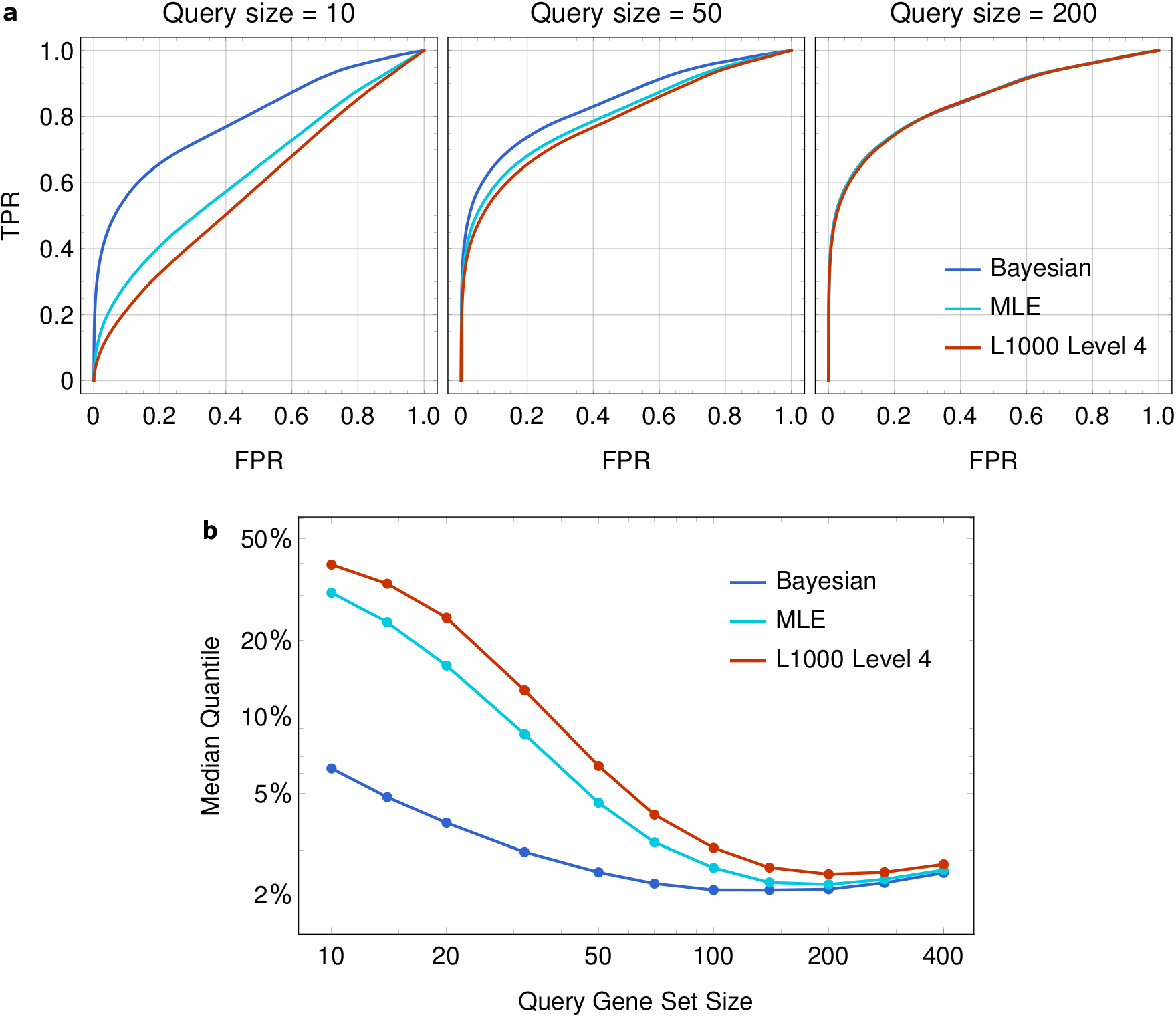
(**a**) Receiver operating characteristic (ROC) curves for replicates identification. Different query size is used for GSEA, and expression profile and features from Bayesian, MLE and L1000 level 4 data are tested. (**b**) The comparison of the median quantile (FPR at TPR = 0.5).

To better illustrate the performance of three methods under different query size, we show the median quantile of GSEA scores between bio replicates,^9^ which is equivalent to the FPR at TPR = 0.5. As shown in Fig. 3(**b**), the median quantile of bio-replicates by all methods become lower as the query size increases up to 200. We notice that the performance of MLE is more similar to that of L1000 than the Bayesian method. As the only difference between Bayesian and MLE is whether peak positions are determined for *z*-score calculation, we conclude that the probabilistic *z*-score inference have a great impact on the performance.

We find that the median quantile by our Bayesian method varies slowly with the query size, while L1000 method have a big improvement when the query size reaches above 50, after which its performance gets close to Bayesian. It indicates that the genes in the query set from our method is more informative when the query size is limited, and the most up/down regulated genes are more stable across bio-replicates by our method. Note that the robustness to small number of query genes is important in real applications. Novel discoveries in pharmacology often involve chemicals and/or cell lines that have not been tested in the L1000 assay, and in such cases, there is no guarantee that a large number of genes can be matched.

### 2.4 Similarity between perturbagens with share targets

L1000 dataset is widely used in drug repurposing and discovery. Signatures in L1000 dataset are compared to each other or target gene sets to estimate the similarity between drugs. To demonstrate the performance in practical problems, we test if our method give similar *z*-score features for similar perturbagens.

Drug target information are obtained from ChEMBL23, and the drugs with the same target annotation are considered similar. In L1000 Phase II dataset, we find 77 groups of similar drugs with an average group size of 2.6 and take them as positive identifications. We use a similar method in measuring the performance as Sec. 2.3 for the combined *z*-scores from Bayesian method, MLE, and L1000 level 5 data. But since the purturbagens are not the same for each cell line in the experiments, we use all the samples as the background instead of separating cell lines.

As shown in Fig. 4(**a**), the Bayesian method performs the best among three methods especially when the query size is small and in the high specificity (low FPR) region. We also show the median quantile of similar drugs with different query size in Fig. 4(**b**). We find that the median quantile from our Bayesian method is lower than the other methods across all query sizes, and the best performance is about 34% when the query set contains about 100 genes.

**Figure 4:**
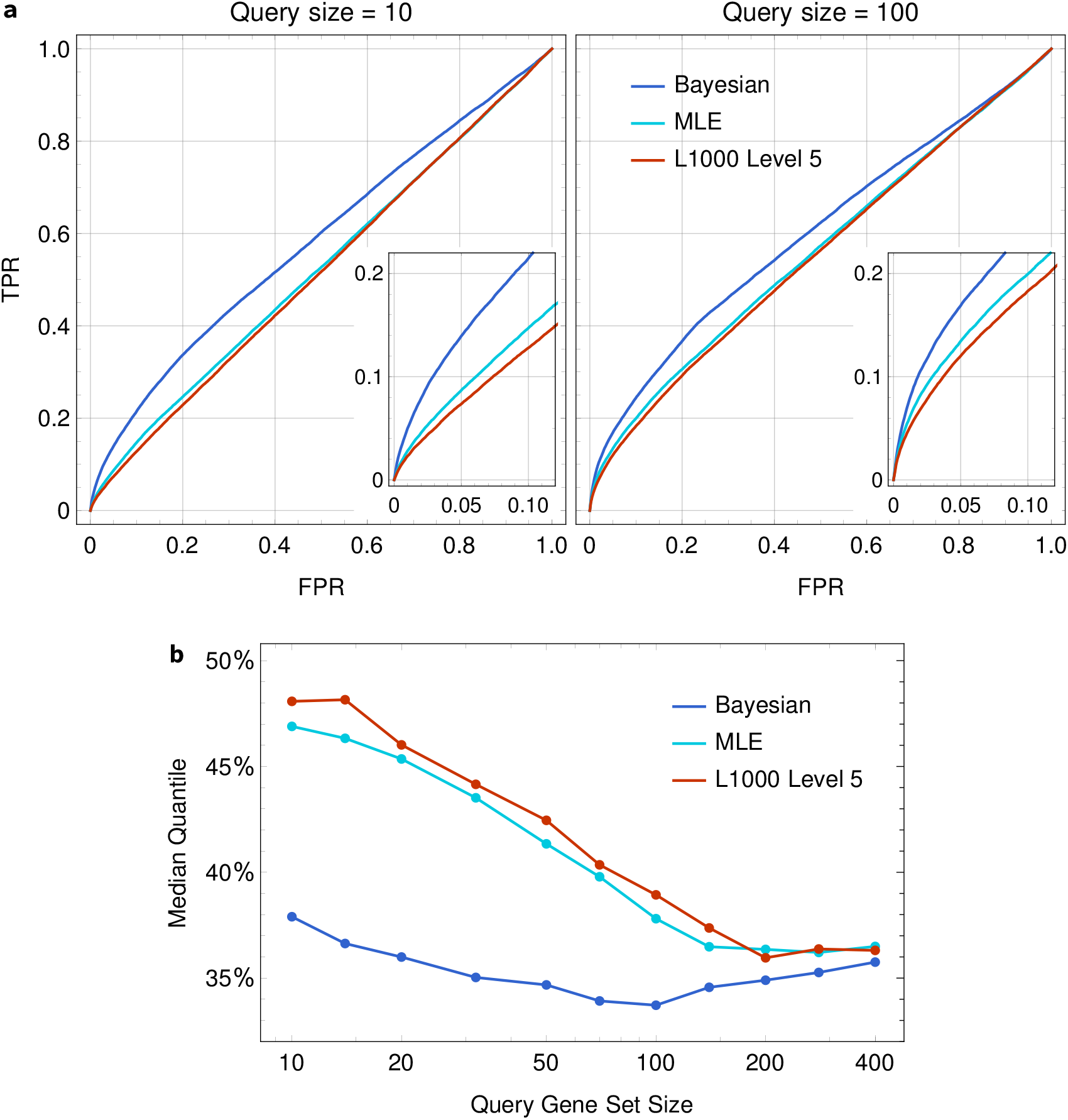
(**a**) ROC curves for similiar perturbagen recognition. Combined *z*-scores from Bayesian, MLE, and L1000 level 5 data are tested. (**b**) The comparison of the median quantile (FPR at TPR = 0.5).

## 3 Discussion

The existence of outliers in FI values are considered a major problem that affects peak detection for the algorithms based on Gaussian models, such as GMM and AGMM. Instead of adding an empirical outlier removal step,^8^ we address this problem by modeling outliers as bead color misidentification. The color of each bead is measured by the Luminex FlexMap 3D platform as three numbers, and there is a small chance that the error in measuring the numbers is large enough for the bead to be identified as a wrong color. We set the rate of misidentification *α*_c_ to be 1% for our precomputed datasets. We notice that the *z*-scores are not sensitive to this rate, where changing *α*_c_ to 3% or 0.3% will affect less than one *z*-score in each profile on average. Additionally, we model the shape of the peaks as Student’s *t*-distributions (see Online Methods), which have a heavier tail than the Gaussian distribution, and the peak locations are less sensitive to outliers compared to Gaussian models.

Peak flip is another problem that is widely discussed.^8, 12^ We address this problem by considering the number of reads in two peaks following a binomial distribution, which is built into our mathematical model. When the number of reads are very close for the two peaks, the likelihood function will show a similar probability for both ways of peak assignment, and the *z*-score from our pipeline is often close to zero which correctly describe the situation. We also note that peak flip is very rare in real data. For instance, a sample with 60 reads and a 2:1 mix ratio, the probability of peak flip is ∼ 0.5%.

As we find the performance of MLE in Fig. 3(**b**) and Fig. 4(**b**) is more similar to that of L1000 rather than Bayesian method, we speculate a possible explanation for this phenomenon as follows. Any deterministic method, however accurate in modeling, picks a precise number for each peak according to some likelihood function. But in the real data, there are many cases where the peak positions are hard to decide. For example, the smaller peak contains too few reads thus hard to be distinguished from noise, or two peaks are not well separated and their locations have a large uncertainty. In those cases, a deterministic method has a small chance to give large *z*-scores to genes that are not regulated. Given that only a little fraction of the genes are actually regulated, the quality of highest and lowest *z*-scores are compromised.

In this paper, we used GSEA as a robust signature comparison method and top up/down regulated genes as the feature for a sample, which is a widely accepted non-parametric way to compare expression profiles. But with changes we made in the signature generation process, different comparison algorithm can be developed to better capture information in signatures.

## 4 Online Methods

### 4.1 Datasets

In this study, we use L1000 small molecule compound data from Broad Institute LINCS L1000 Phase II datasets (https://www.ncbi.nlm.nih.gov/geo/query/acc.cgi?acc=GSE70138), which are categorized into five levels as follows.

Level 1 data contain raw fluorescent intensity (FI) values and three-dimensional color codes for each bead measured by the Luminex FlexMap 3D platform. An FI value is proportional to the transcript abundance of the gene associated to a type of beads. The types of the beads are inferred from their color codes and marked by numbers between 1 and 500 or marked by 0 meaning that they cannot be attributed to any one of the 500 types.

Level 2 data contain gene expression values for the 978 landmark genes, 976 of which are grouped into pairs and associated with 488 types of beads and the rest two of which are associated with two separate types of beads. A peak deconvolution process^2^ is employed to measure one or two expression values from the FI values for each type of beads in each well. A profile consisting 978 expressions values is therefore obtained.

Level 3 data contain normalized gene expression values. The normalization process is divided into two parts: L1000 invariant set scaling (LISS) and quantile normalization (QNORM). In LISS, all gene expression values in a well, i.e. a sample, are scaled to a set of pre-defined control genes. Then, the expression values are quantile normalized across all wells on each plate.

Level 4 data contain the *z*-scores for each gene with all expression values of that gene on a plate as the background. *z*-scores indicate the levels at which genes are differentially expressed. They are then combined across biological replicates to obtain level 5 data.

### 4.2 L1000 invariant set scaling and quality control

Since the amplification factor of each sample is different, L1000 added 80 control transcripts to each well, whose expression levels are empirically found to be invariant as calibration set. These genes are grouped into 10 levels with 8 each so that the median expression of each level should follow a similar increasing trend. The median expression levels in terms their fluorescent intensities are then compared with reference values and a relation between them can be inferred by fitting various template functions on a logarithmic scale.^10^

L1000 used a power law relation *y* = *ax*^*b*^ + *c* in their pipline, where *x* is the unscaled data and *y* the scaled data.^2^ In this study, we employ a two-step linear scaling as follows. First, we rescale the median expressions of 10 invariant sets by fitting them against the reference values linearly, and we obtain the averages and standard deviations of those scaled expressions within each perturbagen group. The resulting calibrated reference expressions shown in Fig. 5. Second, we fit the median expressions of the invariant sets against the calibrated expressions linearly and use this relation to rescale the expressions of the landmark genes.

**Figure 5:**
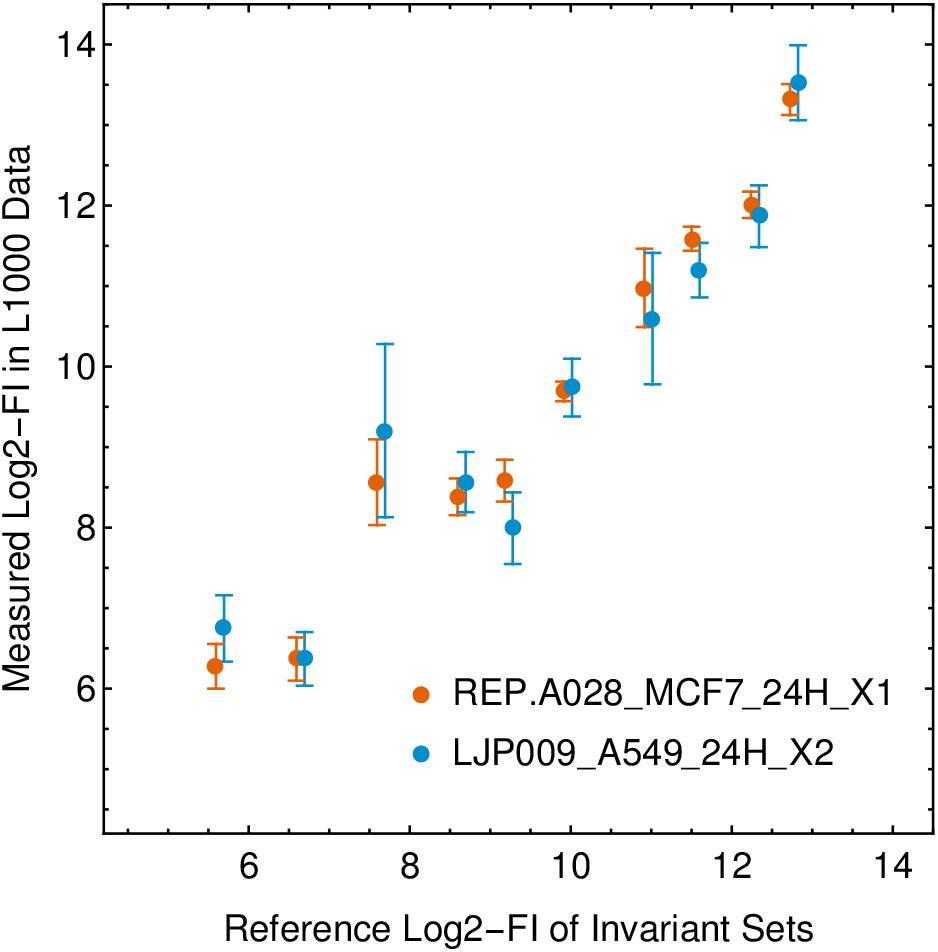
The calibrated reference expressions values of the invariant set compared with the original ones. The error bars show the ranges corresponding to ±1 standard deviation. The data points are horizontally offset for better visibility.

The two-step linear scaling have two advantages. First, since the median expressions of the invariant sets vary across plates (see Fig. 5), calibrated reference expressions depending on the purturbagen groups are needed capture this variability. Second, the linear scaling in the second step depends on only two fitted parameters, one less than the power law scaling.

We identify low-quality samples according to two parameters: the goodness of fit in terms of *χ*^2^ and the slope *a* of the relation, where an exceptionally large slope suggests a failed amplification. Fig. 6 shows the distribution of these two parameters. L1000 level 1 data have 4.0% of the samples missing due to its quality control and we remove additional 3.0% of the samples with *a* > 3.0 or *χ*^2^ > 4.0 as our quality control.

**Figure 6:**
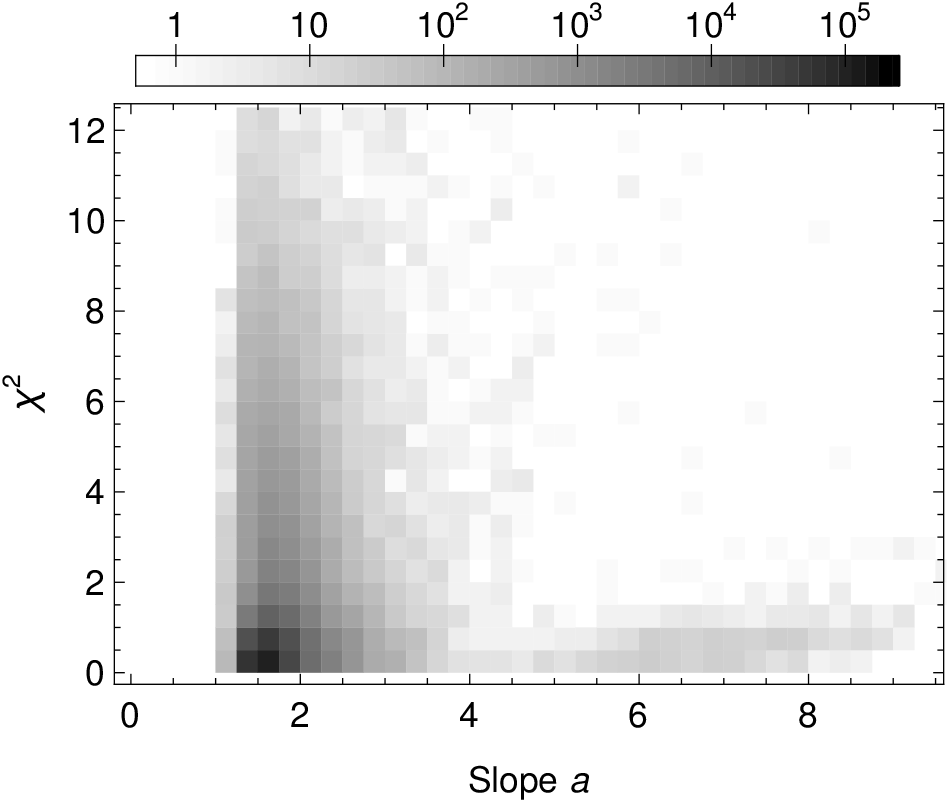
The distribution of the slope and *χ*^2^ of the fitted relation between unscaled expression values and calibrated reference values.

### 4.3 Peak deconvolution

The Luminex FlexMap 3D platform can detect 500 different bead colors. In order to measure 978 transcripts in a sample within a single batch, almost all of them are grouped into pairs. For each pair, barcodes of two transcripts are coupled to beads with the same color and they are mixed in a ratio of 2:1. Therefore, the distributions of reads from these bead colors will form two peaks, whose expression levels should be measured by the peak deconvolution algorithm. In the design of L1000 system, the pairing of genes is optimized in order to minimize the confusion in peak deconvolution, but we note that it is inevitable that the peaks in some of the distributions are hard to be differentiated.

In this study, we develop a new peak deconvolution algorithm based on Bayesian probability model. L1000 samples typically have dozens of reads per bead color in each sample. In our model, each of these reads will either come from one of the genes that are coupled to that bead color or come from an arbitrary bead in the same sample due to color misidentification with a small rate *α*_c_ ~ 1%. Suppose that *N* measurements are made for a specific color; we can derive the probability of the number of measurements that are associated with both genes *N*_hi_, *N*_lo_ and the number of color misidentifications, or the background *N*_bg_, where *N* = *N*_hi_ + *N*_lo_ + *N*_bg_. Given that *α*_c_ ≪ 1, we assume that *N*_bg_ follows the Poisson distribution with *λ* = *α*_c_*N* and *N*_hi_ follows the binomial distribution *B*(*N* − *N*_bg_, 2/3).

The shapes of the peaks reflect the uncertainty of Luminex intensity measurements, and they should depend on their expression levels only. Using L1000 level 2 data as a reference, we first filter the profiles of reads that have well separated peaks, where the distance between the center of two peaks is at least 3 in terms of log2 expression. Then, we pick discrete expression levels between 4 and 12, extract the profiles that have one of their peaks located in the neighborhood of each of them, and take the average profiles as the shapes of the peaks. We use Student’s *t*-distribution to model the peaks. As is shown in Fig. 7, a wide range of degrees of freedom (DOF) values can describe the peaks across all expression levels, thus we choose a fixed DOF of three. We have also measured an empirical relation between the scale parameter *σ* and the log2 expression value *x*, shown in Fig. 8, which can be fitted by an analytical expression as

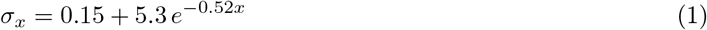

with a mean absolute error of 0.009. The shape of a single peak centered at log2 expression level *x* is therefore given by

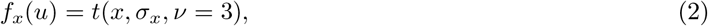

where *u* is the log2-FI. As for bead color misidentification, we assume that the distribution of these reads follows the distribution of all reads *f*_bg_(*u*) from the same well.

**Figure 7:**
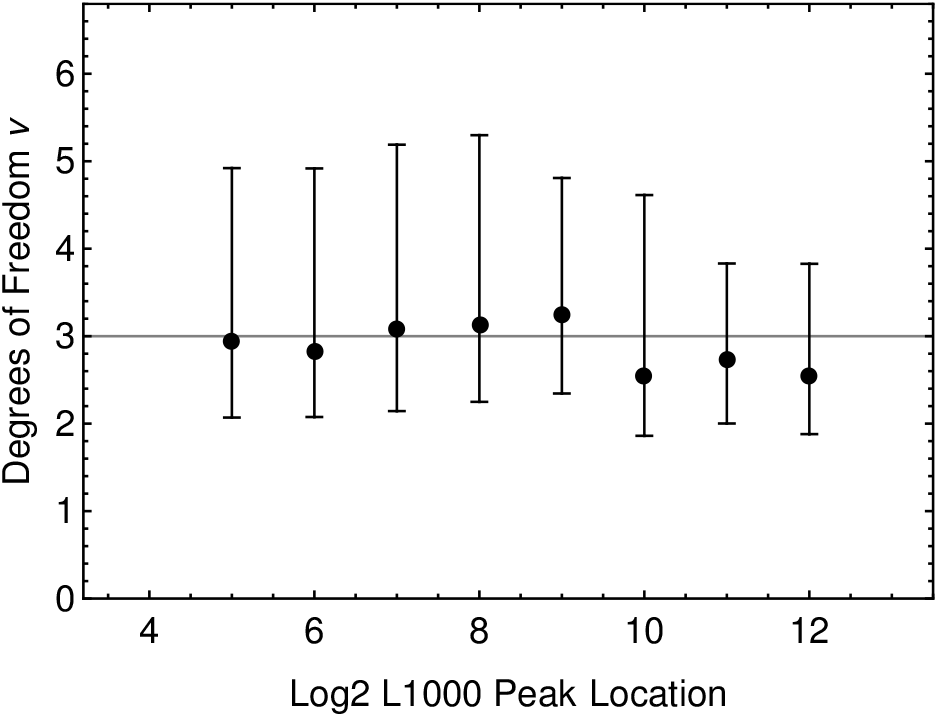
The median and 68% confidence interval of the best fit degrees of freedom of Student’s *t*-distribution. Our choice of three degrees of freedom is shown by a gray line.

**Figure 8:**
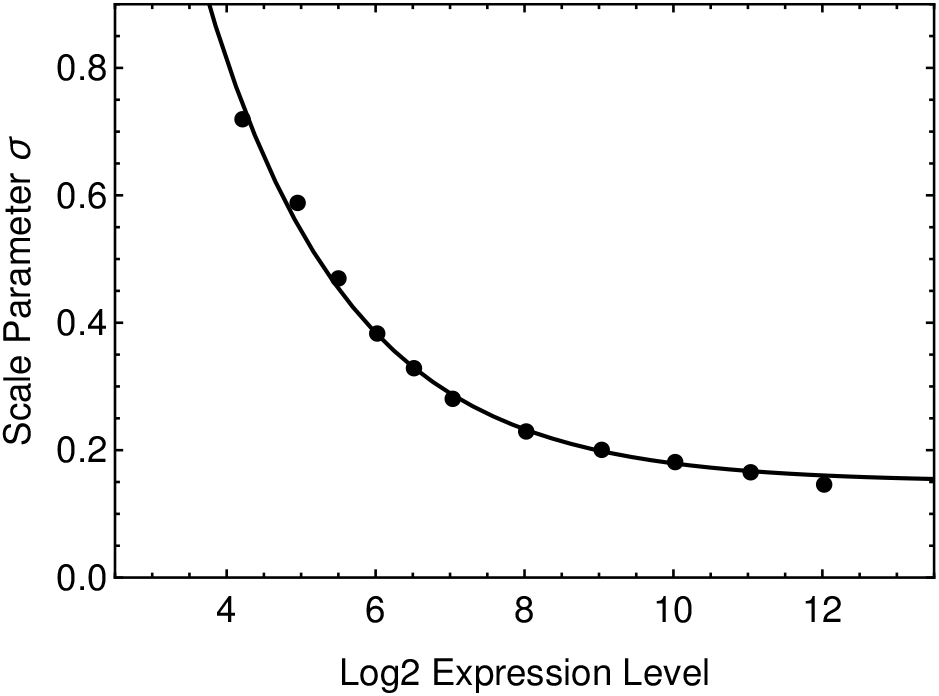
The relation between the scale parameter of the Student’s *t*-distribution and the log2 expression level, i.e. the center of an isolated peak.

With the shape of peaks and background known, we can determine the probability of a read to be found with fluorescent intensity *u*_*i*_ described as a summation of probability for it to come from either of the two genes or background:

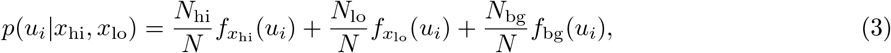

where *x*_hi_ and *x*_lo_ are the center of the two peaks. The posterior distribution of *x*_hi_ and *x*_lo_ is given by

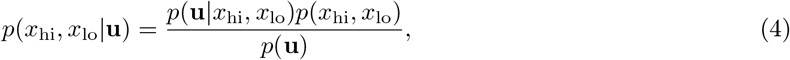

where **u** denotes all of the measurements *u*_*i*_ (*i* = 1, 2,…, *N*). We adopt a uniform prior on *x*_hi_ and *x*_lo_, and the posterior distribution becomes

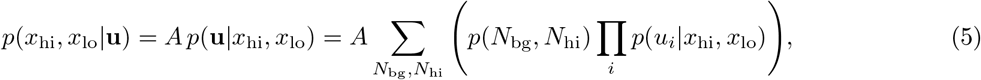

where *A* denotes a normalizing constant. We note that the calculation of the likelihood function is not trivial, and we show the simplifications of the function in Supplementary Materials.

In order to simplify further analysis, we marginalize *x*_hi_ and *x*_lo_ in all samples to get *p*(*x*_*g*_|**u**), or simply *p*(*x*_*g*_), where *g* = 1, 2, …, 978 are the indices of the genes. Compared to L1000 level 2 data, in which *x*_*g*_ have precise values, our algorithm of peak deconvolution gives two probability distributions, revealing the uncertainty of these estimations.

### 4.4 Quantile normalization

Quantile normalization (QNORM) standardize the shape of expression profile distributions among the wells on each plate. In L1000 level 3 data, QNORM is done by first sorting the expression levels within samples, then setting the *i*-th highest values in each sample by their median value.

In this study, we use a similar way to do quantile normalization. First, we add together all the marginal distributions of all genes in each sample to get the overall distribution of expression levels

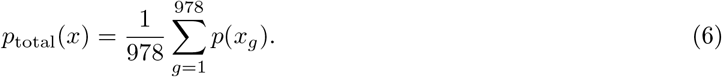

Then, we define the relative fluorescent intensity *r* as

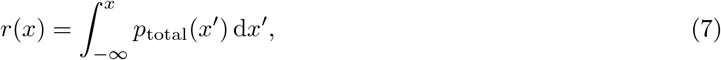

which has a value in the interval of [0, 1]. Finally, we standardize *p*_total_(*x*) as a uniform distribution *U* (0, 1) to get the quantile normalized distribution of individual expression levels as

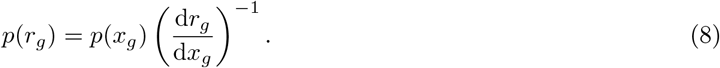

### 4.5 *z*-score inference

With QNORM done, the profiles of different samples are now comparable to each other. L1000 uses *z*-score to represent relative gene expression. In a normal distributed population, *z*-score is defined by *z*_*g*_ = (*x*_*g*_ − *µ*_*g*_)/*σ*_*g*_, where *µ*_*g*_ denotes the average value of the population and *σ*_*g*_ the standard deviation. Due to the fact that the distributions of gene expression typically have heavier tails, i.e. having more extreme values than the normal distribution, L1000 uses the median and median absolute difference (MAD) instead and defines the *z*-scores by

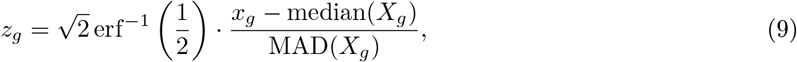

which is equivalent to the original one in the case of normal distribution. A common choice of the population *X*_*g*_ is the expression levels *x*_*g*_ of all samples on a plate.

In our case, since we depict the expression level by a probability distribution instead of a number, both definitions of the *z*-score cannot be used directly. However, we note that the *z*-score, assuming a normal distribution, is related to the quantile of the expression level in the population by

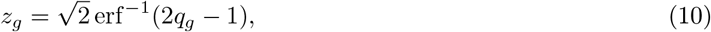

where the expected value of the quantile *q*_*g*_ is the probability that the expression level

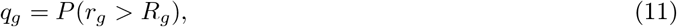

in which the probability distribution of *R*_*g*_ equals to the summation of those of all samples on a plate

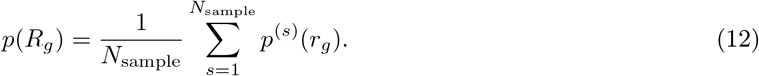

### 4.6 Combining replicates

We follow the L1000 pipeline and use a weighed average in combining replicates based on the correlations among the bio-replicates. Denote the combined *z*-scores of all genes as **z**_c_ and the *z*-scores of the *i*-th bio-replicate as **z**^(*i*)^, we have

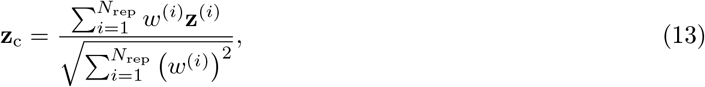

where *N*_rep_ is the number of replicates and the weights *w*^(*i*)^ are defined as

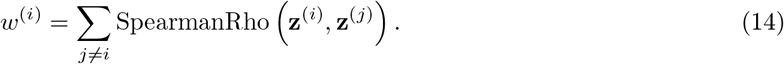

## Acknowledgments

This work was supported by Grant Number R01LM011986 from the National Library of Medicine (NLM), Grant Number R01GM122845 from the National Institute of General Medical Sciences (NIGMS), and Grand Number R01AD057555 of National Institute of Aging of the National Institute of Health (NIH) as well as CUNY High Performance Computing Center.

## Author contributions

Y.Q., T.L. developed and implemented methods, designed the experiments, performed experiments, analyzed the data, and wrote the manuscript; H.L. provided and analyzed data; L.X. conceived the project, designed the experiments, analyzed the data, and wrote the manuscript.

