## Supplementary information for "A Bayesian approach to accurate and robust signature detection on LINCS L1000 data"

### Supplementary A The calculation of the likelihood function

In the main article, the likelihood function of the peak locations are given by

$$p(x_{\rm hi}, x_{\rm lo}|\mathbf{u}) = A\, p(\mathbf{u}|x_{\rm hi}, x_{\rm lo}) = A \sum_{N_{\rm bg}, N_{\rm hi}} \left( p(N_{\rm bg}, N_{\rm hi}) \prod_i p(u_i|x_{\rm hi}, x_{\rm lo}) \right), \tag{1}$$

but we note that it is not trivial to do such a computation. We calculate each likelihood function on a $400 \times 400$ grid for all combinations of $x_{\rm hi}$ and $x_{\rm lo}$, and the total number of operations for a single sample is expected to be $\sim 10^{13}$. Since L1000 dataset contain $\sim 10^6$ samples, simplifications are necessary to carry out the computation. In this section, we introduce several simplifications to $p(\mathbf{u}|x_{\rm hi}, x_{\rm lo})$, which is equivalent to the likelihood function $p(x_{\rm hi}, x_{\rm lo}|\mathbf{u})$ as has been shown in the main article.

Note that the chance of bead color misidentification is small $\alpha_{\rm c} \sim 1\%$, so the probability of having a large $N_{\rm bg}$ is negligible. L1000 typically have 50 reads for each bead color in one sample, which means $\langle N_{\rm bg} \rangle \approx 0.5$, and probability of $0 \le N_{\rm bg} \le 3$ is around 99.8%. Therefore, we isolate $N_{\rm bg}$ from the summation and only do the summation over four different $N_{\rm bg}$ with the largest probabilities:

$$p(\mathbf{u}|x_{\rm hi}, x_{\rm lo}) = \sum_{N_{\rm bg}} p(N_{\rm bg}) \sum_{N_{\rm hi}} \left( p(N_{\rm hi}|N_{\rm bg}) \prod_i p(u_i|x_{\rm hi}, x_{\rm lo}) \right). \tag{2}$$

Now, we simplify the expression in the summation over $N_{\rm hi}$ in Eqn.(2):

$$L(N_{\rm hi}) \equiv p(N_{\rm hi}|N_{\rm bg}) \prod_i p(u_i|x_{\rm hi}, x_{\rm lo}). \tag{3}$$

The first part is the conditional probability $p(N_{\rm hi}|N_{\rm bg})$. It follows the binomial distribution $B(N - N_{\rm bg}, 2/3)$, and it can be approximated by a normal distribution with the same mean and variance:

$$N_{\rm hi} \sim \mathcal{N}\left( \mu_{\rm hi} = \frac{2}{3}(N - N_{\rm bg}), \sigma_{\rm hi}^2 = \frac{2}{9}(N - N_{\rm bg}) \right), \tag{4}$$

so

$$p(N_{\rm hi}|N_{\rm bg}) = \frac{1}{\sqrt{2\pi\sigma_{\rm hi}^2}} \exp\left( -\frac{(N_{\rm hi} - \mu_{\rm hi})^2}{2\sigma_{\rm hi}^2} \right). \tag{5}$$

The second part is the conditional probability $p(u_i|x_{\rm hi}, x_{\rm lo})$. When $x_{\rm hi}$, $x_{\rm lo}$, and $N_{\rm bg}$ are fixed, by Eqn.(**??**), $p(u_i|x_{\rm hi}, x_{\rm lo})$ linearly depends on $N_{\rm hi}$ given $u_i$. Hence, we can express $p(u_i|x_{\rm hi}, x_{\rm lo})$ in the following form:

$$p(u_i|x_{\rm hi}, x_{\rm lo}) = \tilde{p}(u_i) + (N_{\rm hi} - \mu_{\rm hi}) \frac{\delta p(u_i)}{\delta N_{\rm hi}}, \tag{6}$$

where the reference point of probability $\tilde{p}(u_i)$ is taken at $N_{\rm hi} = \mu_{\rm hi}$

$$\tilde{p}(u_i) = \left(1 - \frac{N_{\rm bg}}{N}\right) \left( \frac{2}{3} f_{x_{\rm hi}}(u_i) + \frac{1}{3} f_{x_{\rm lo}}(u_i) \right) + \frac{N_{\rm bg}}{N} f_{\rm bg}(u_i), \tag{7}$$

and change in the probability for incremental $N_{\rm hi}$ is

$$\frac{\delta p(u_i)}{\delta N_{\rm hi}} = \frac{f_{x_{\rm hi}}(u_i) - f_{x_{\rm lo}}(u_i)}{N}. \tag{8}$$

The approximated $p(N_{\rm hi}|N_{\rm bg})$ and the new form of $p(u_i|x_{\rm hi}, x_{\rm lo})$ allow us to rewrite Eqn.(3) as

$$\begin{aligned}
\log L(N_{\rm hi}) &= \log\left( p(N_{\rm hi}|N_{\rm bg}) \prod_i p(u_i|x_{\rm hi}, x_{\rm lo}) \right) && (9) \\
&= \log p(N_{\rm hi}|N_{\rm bg}) + \sum_i \log p(u_i|x_{\rm hi}, x_{\rm lo}) && (10) \\
&= -\frac{1}{2}\log\left(2\pi\sigma_{\rm hi}^2\right) - \frac{(N_{\rm hi} - \mu_{\rm hi})^2}{2\sigma_{\rm hi}^2} + \sum_i \log p(u_i|x_{\rm hi}, x_{\rm lo}) && (11)
\end{aligned}$$

Note that the first two terms in Eqn.(9) constitute a second order polynomial with respect to $(N_{\rm hi} - \mu_{\rm hi})$. Thus, it is useful to expand $\log p(u_i|x_{\rm hi}, x_{\rm lo})$ to the second order centered at $N_{\rm hi} = \mu_{\rm hi}$ as

$$\begin{aligned} \log p(u_i|x_{\rm hi}, x_{\rm lo}) &= \log\left(\tilde{p}(u_i) + (N_{\rm hi} - \mu_{\rm hi})\frac{\delta p(u_i)}{\delta N_{\rm hi}}\right) && (12) \\ &\approx \log\tilde{p}(u_i) + \frac{(N_{\rm hi} - \mu_{\rm hi})}{\tilde{p}(u_i)}\frac{\delta p(u_i)}{\delta N_{\rm hi}} - \frac{1}{2}\left(\frac{(N_{\rm hi} - \mu_{\rm hi})}{\tilde{p}(u_i)}\frac{\delta p(u_i)}{\delta N_{\rm hi}}\right)^2, && (13) \end{aligned}$$

so that $\log L(N_{\rm hi})$ can be expressed fully as a second order polynomial as

$$\log L(N_{\rm hi}) = a + b(N_{\rm hi} - \mu_{\rm hi}) - c(N_{\rm hi} - \mu_{\rm hi})^2, \quad (14)$$

where the coefficients are

$$\begin{aligned} a &= -\frac{1}{2}\log\left(2\pi\sigma_{\rm hi}^2\right) + \sum_i \log\tilde{p}(u_i), && (15) \\ b &= \sum_i \frac{1}{\tilde{p}(u_i)}\frac{\delta p(u_i)}{\delta N_{\rm hi}}, && (16) \\ c &= \frac{1}{2\sigma_{\rm hi}^2} + \frac{1}{2}\sum_i\left(\frac{1}{\tilde{p}(u_i)}\frac{\delta p(u_i)}{\delta N_{\rm hi}}\right)^2. && (17) \end{aligned}$$

With the above transformations, the summation over $N_{\rm hi}$ in Eqn.(2) can be approximated as a definite integral over $N_{\rm hi}$, and finally, the likelihood function can be calculated by

$$\begin{aligned} p(\mathbf{u}|x_{\rm hi}, x_{\rm lo}) &= \sum_{N_{\rm bg}} p(N_{\rm bg}) \sum_{N_{\rm hi}} L(N_{\rm hi}) && (18) \\ &= \sum_{N_{\rm bg}} p(N_{\rm bg}) \sum_{N_{\rm hi}} \exp\left(a + b(N_{\rm hi} - \mu_{\rm hi}) - c(N_{\rm hi} - \mu_{\rm hi})^2\right) && (19) \\ &\approx \sum_{N_{\rm bg}} p(N_{\rm bg}) \int_{N_{\rm hi}} {\rm d}N_{\rm hi}\, \exp\left(a + b(N_{\rm hi} - \mu_{\rm hi}) - c(N_{\rm hi} - \mu_{\rm hi})^2\right) && (20) \\ &= \sum_{N_{\rm bg}} p(N_{\rm bg}) \sqrt{\frac{\pi}{c}}\, \exp\left(a + \frac{b^2}{4c}\right), && (21) \end{aligned}$$

where the parameters $a$, $b$, and $c$ depend on $\mathbf{u}$ and $N_{\rm bg}$ implicitly.

#### Supplementary B Peak locations for a sample well by Bayesian, MLE, and L1000

Here we show the full list of peak locations in well REP.A028_MCF7_24H_X2_B25_D11 given by our Bayesian method, Bayesian (MLE), and L1000 level 2 data. The results from L1000 peak deconvolution and AGMM are shown in red and green arrows, where the thick and thin arrows indicate the peaks with high (2/3) and low (1/3) abundances respectively. The results from our method are shown as probability distributions for high and low peaks in thick and thin blue curves respectively.

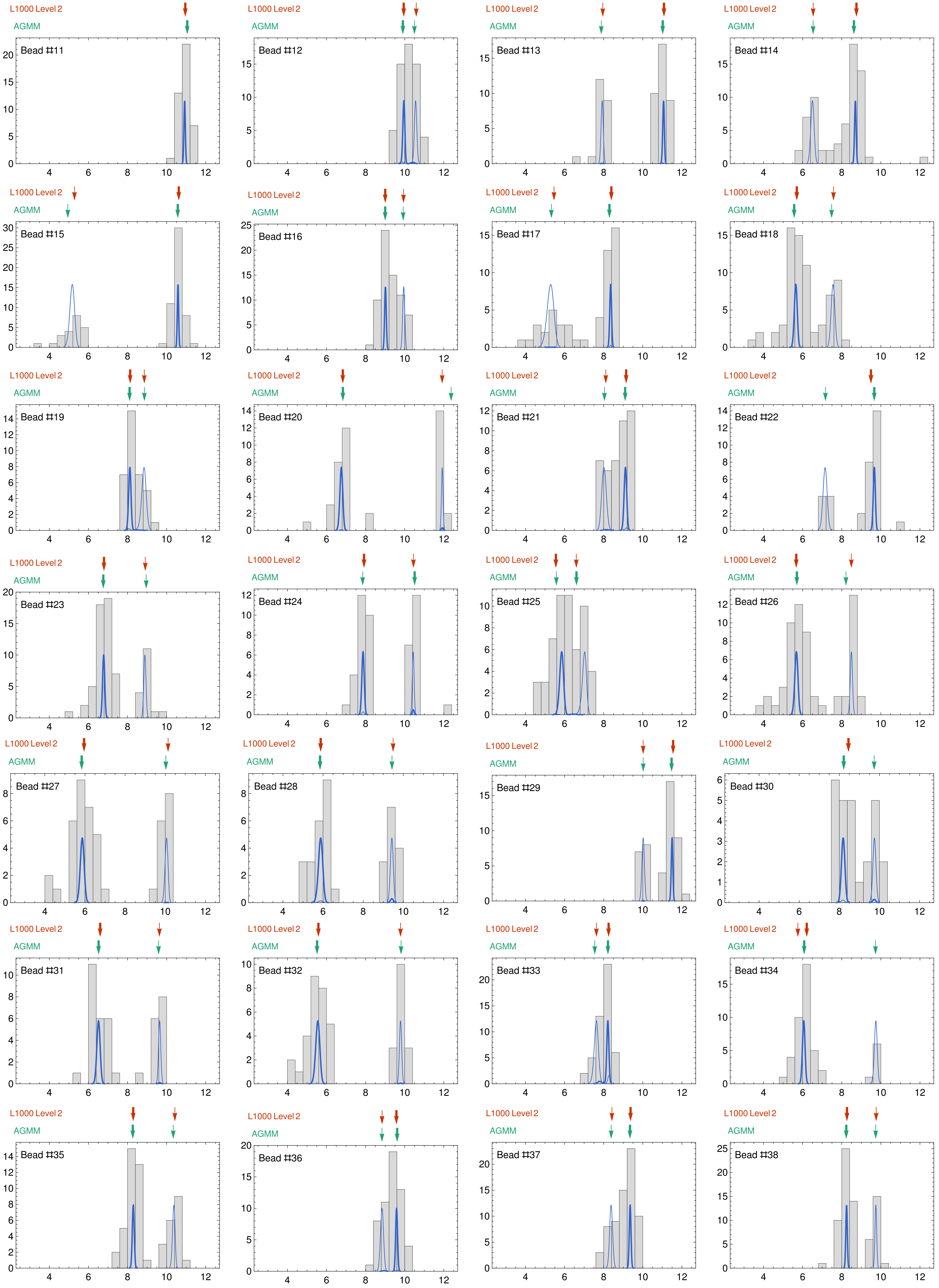

L1000 Level 2
AGMM
Bead #11
Bead #12
Bead #13
Bead #14
Bead #15
Bead #16
Bead #17
Bead #18
Bead #19
Bead #20
Bead #21
Bead #22
Bead #23
Bead #24
Bead #25
Bead #26
Bead #27
Bead #28
Bead #29
Bead #30
Bead #31
Bead #32
Bead #33
Bead #34
Bead #35
Bead #36
Bead #37
Bead #38

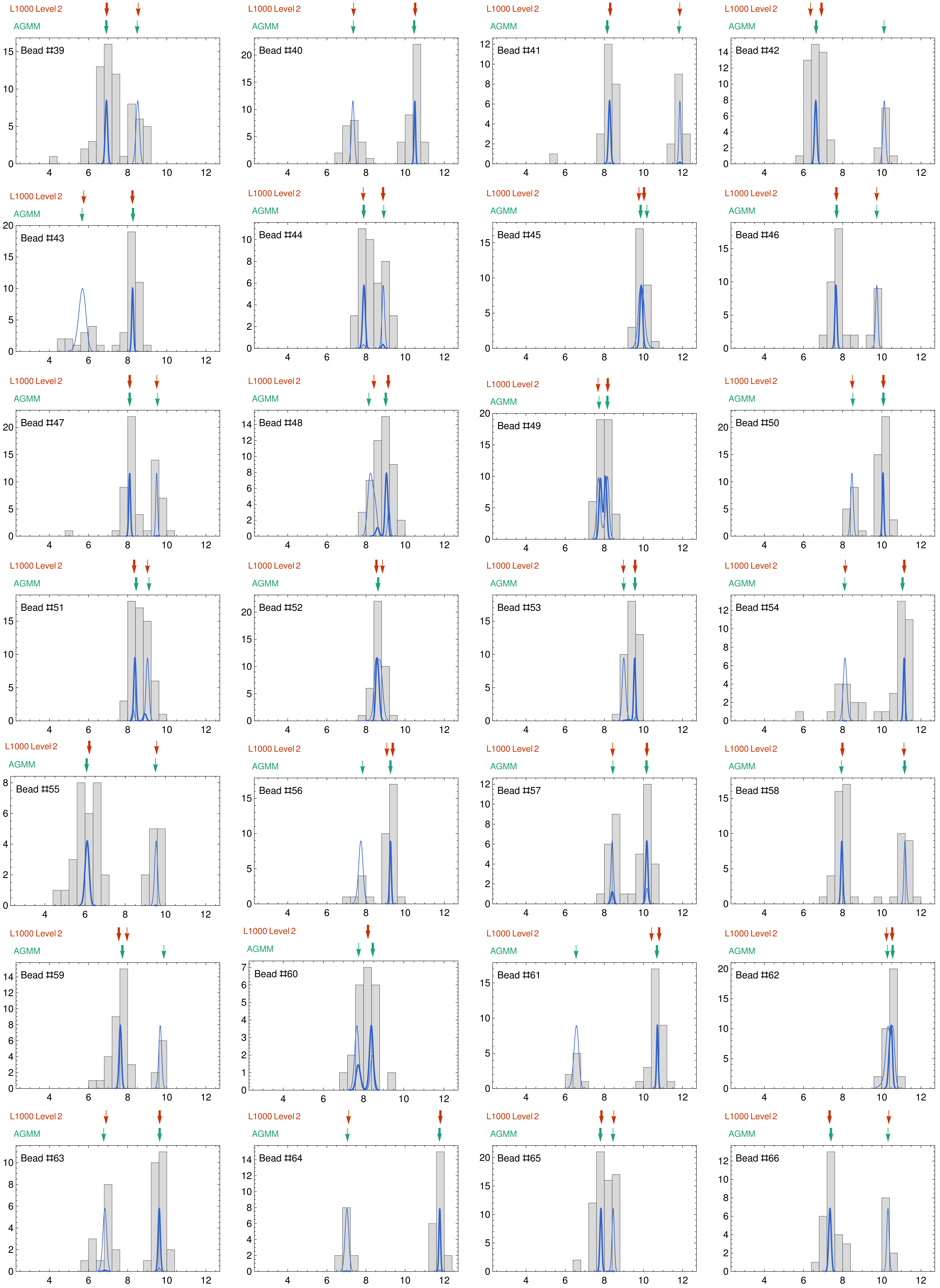
L1000 Level 2
AGMM
Bead #39
Bead #40
Bead #41
Bead #42
Bead #43
Bead #44
Bead #45
Bead #46
Bead #47
Bead #48
Bead #49
Bead #50
Bead #51
Bead #52
Bead #53
Bead #54
Bead #55
Bead #56
Bead #57
Bead #58
Bead #59
Bead #60
Bead #61
Bead #62
Bead #63
Bead #64
Bead #65
Bead #66

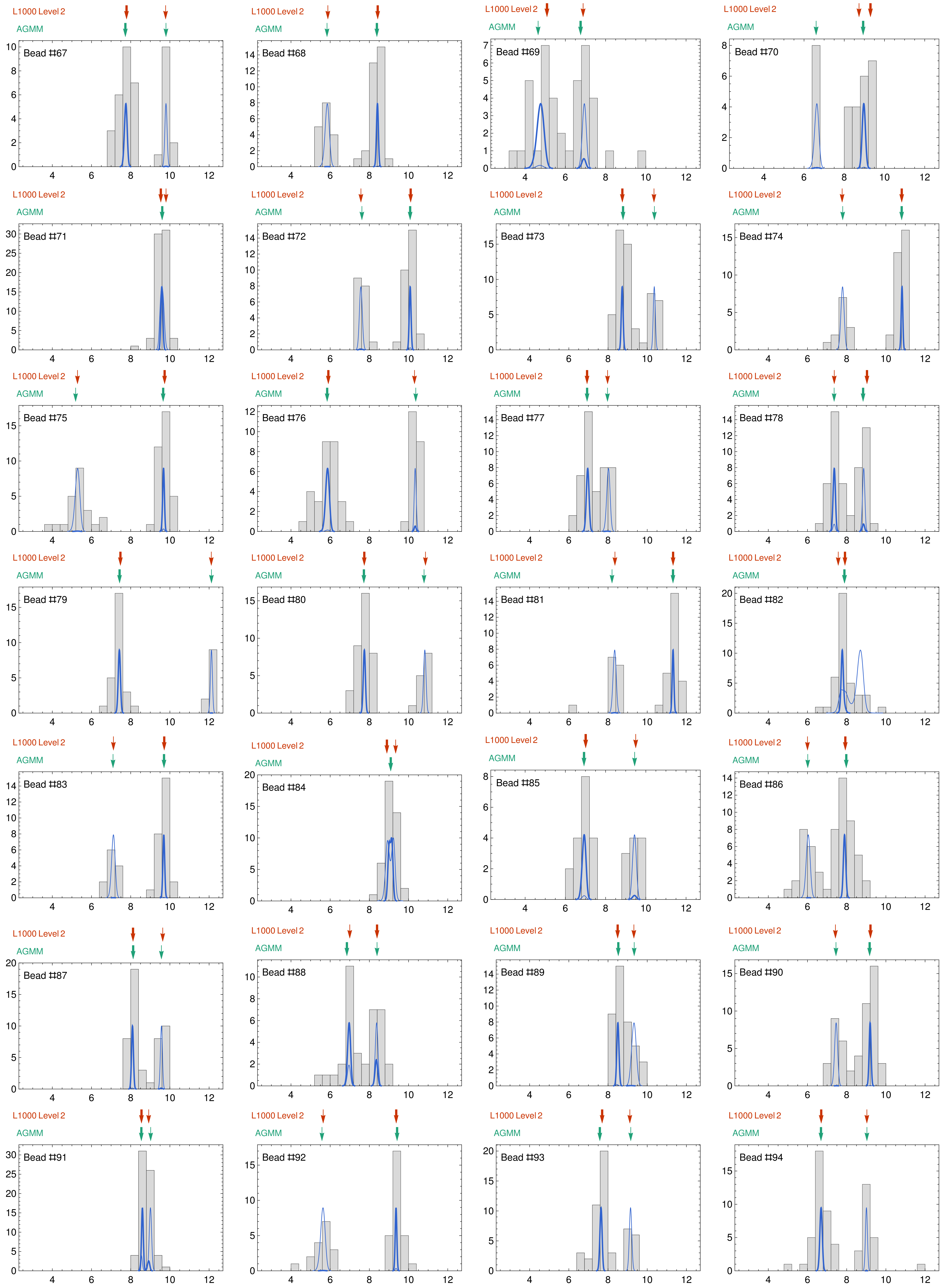
L1000 Level 2
AGMM
Bead #67
Bead #68
Bead #69
Bead #70
Bead #71
Bead #72
Bead #73
Bead #74
Bead #75
Bead #76
Bead #77
Bead #78
Bead #79
Bead #80
Bead #81
Bead #82
Bead #83
Bead #84
Bead #85
Bead #86
Bead #87
Bead #88
Bead #89
Bead #90
Bead #91
Bead #92
Bead #93
Bead #94

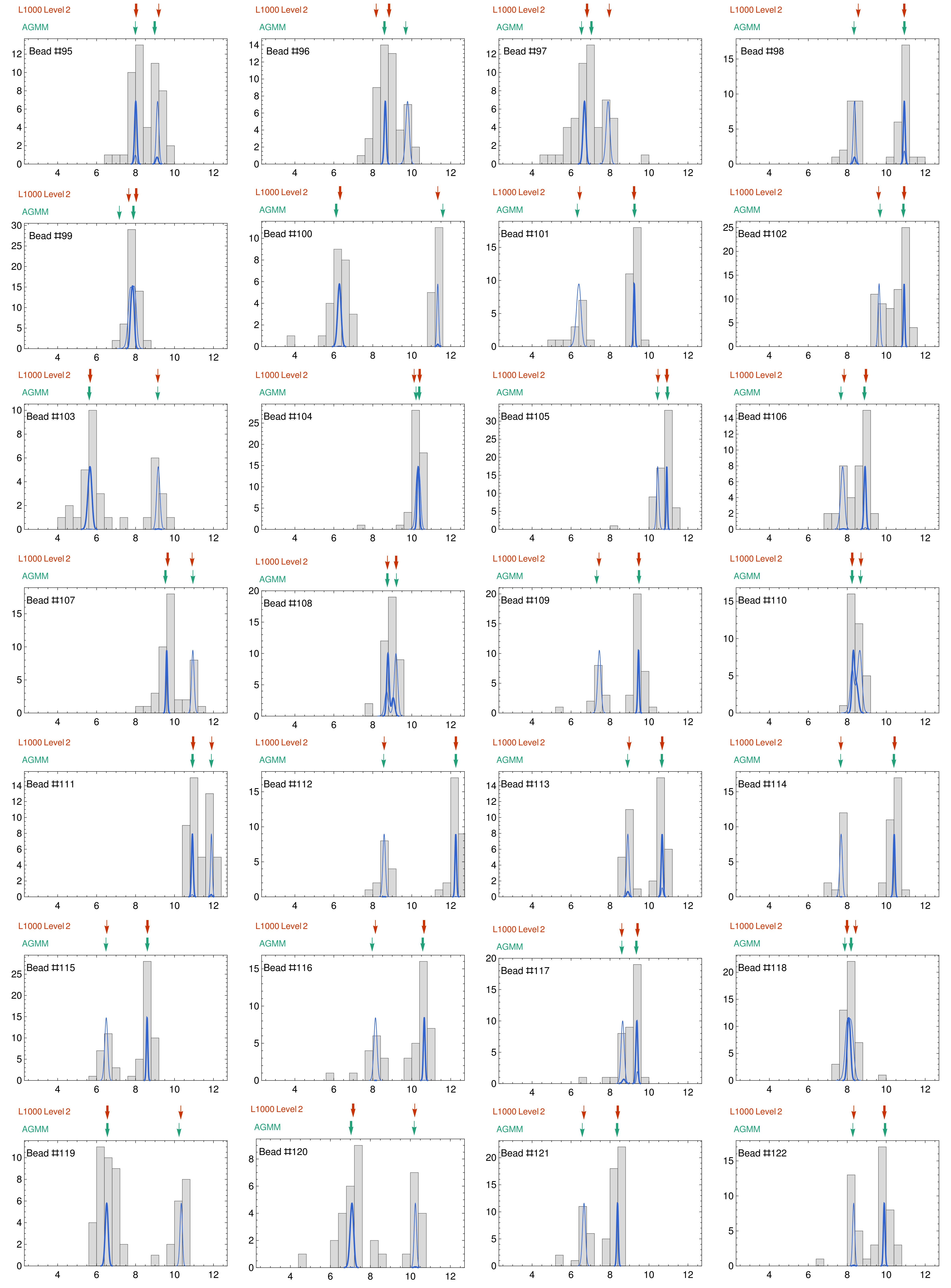

L1000 Level 2
AGMM
Bead #95
Bead #96
Bead #97
Bead #98
Bead #99
Bead #100
Bead #101
Bead #102
Bead #103
Bead #104
Bead #105
Bead #106
Bead #107
Bead #108
Bead #109
Bead #110
Bead #111
Bead #112
Bead #113
Bead #114
Bead #115
Bead #116
Bead #117
Bead #118
Bead #119
Bead #120
Bead #121
Bead #122

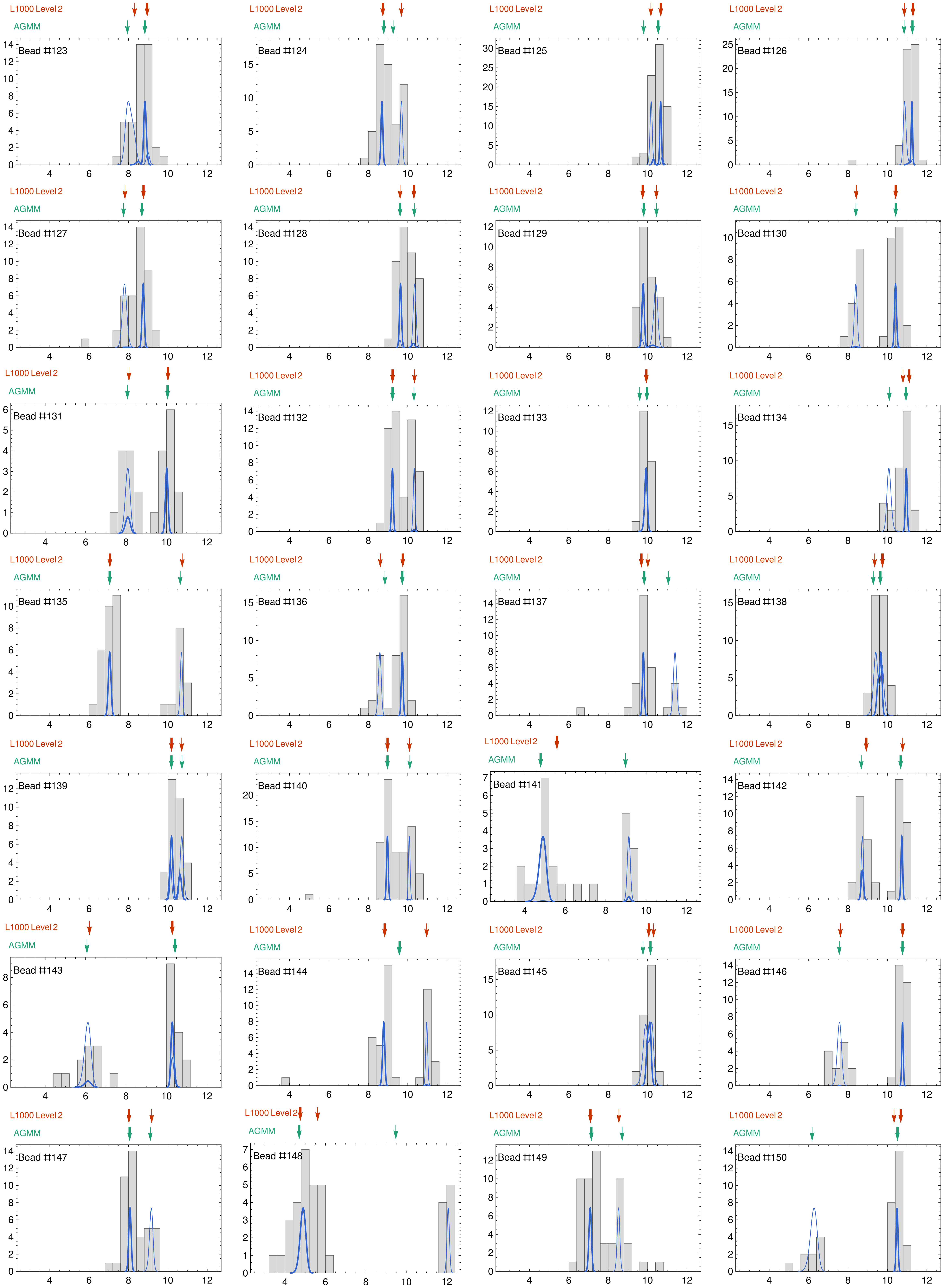

L1000 Level 2
AGMM
Bead #123
Bead #124
Bead #125
Bead #126
Bead #127
Bead #128
Bead #129
Bead #130
Bead #131
Bead #132
Bead #133
Bead #134
Bead #135
Bead #136
Bead #137
Bead #138
Bead #139
Bead #140
Bead #141
Bead #142
Bead #143
Bead #144
Bead #145
Bead #146
Bead #147
Bead #148
Bead #149
Bead #150

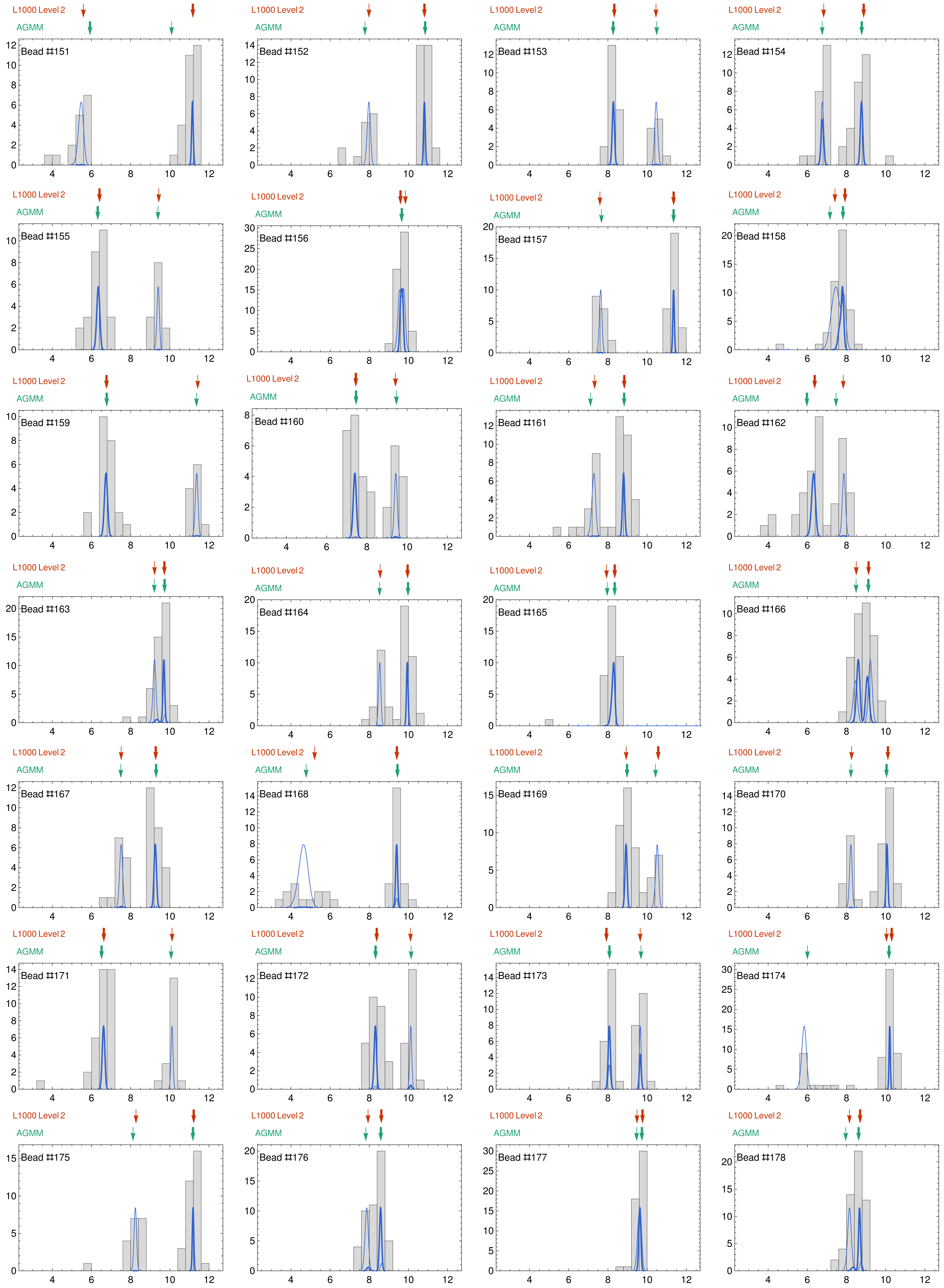

L1000 Level 2
AGMM
Bead #151
L1000 Level 2
AGMM
Bead #152
L1000 Level 2
AGMM
Bead #153
L1000 Level 2
AGMM
Bead #154
L1000 Level 2
AGMM
Bead #155
L1000 Level 2
AGMM
Bead #156
L1000 Level 2
AGMM
Bead #157
L1000 Level 2
AGMM
Bead #158
L1000 Level 2
AGMM
Bead #159
L1000 Level 2
AGMM
Bead #160
L1000 Level 2
AGMM
Bead #161
L1000 Level 2
AGMM
Bead #162
L1000 Level 2
AGMM
Bead #163
L1000 Level 2
AGMM
Bead #164
L1000 Level 2
AGMM
Bead #165
L1000 Level 2
AGMM
Bead #166
L1000 Level 2
AGMM
Bead #167
L1000 Level 2
AGMM
Bead #168
L1000 Level 2
AGMM
Bead #169
L1000 Level 2
AGMM
Bead #170
L1000 Level 2
AGMM
Bead #171
L1000 Level 2
AGMM
Bead #172
L1000 Level 2
AGMM
Bead #173
L1000 Level 2
AGMM
Bead #174
L1000 Level 2
AGMM
Bead #175
L1000 Level 2
AGMM
Bead #176
L1000 Level 2
AGMM
Bead #177
L1000 Level 2
AGMM
Bead #178

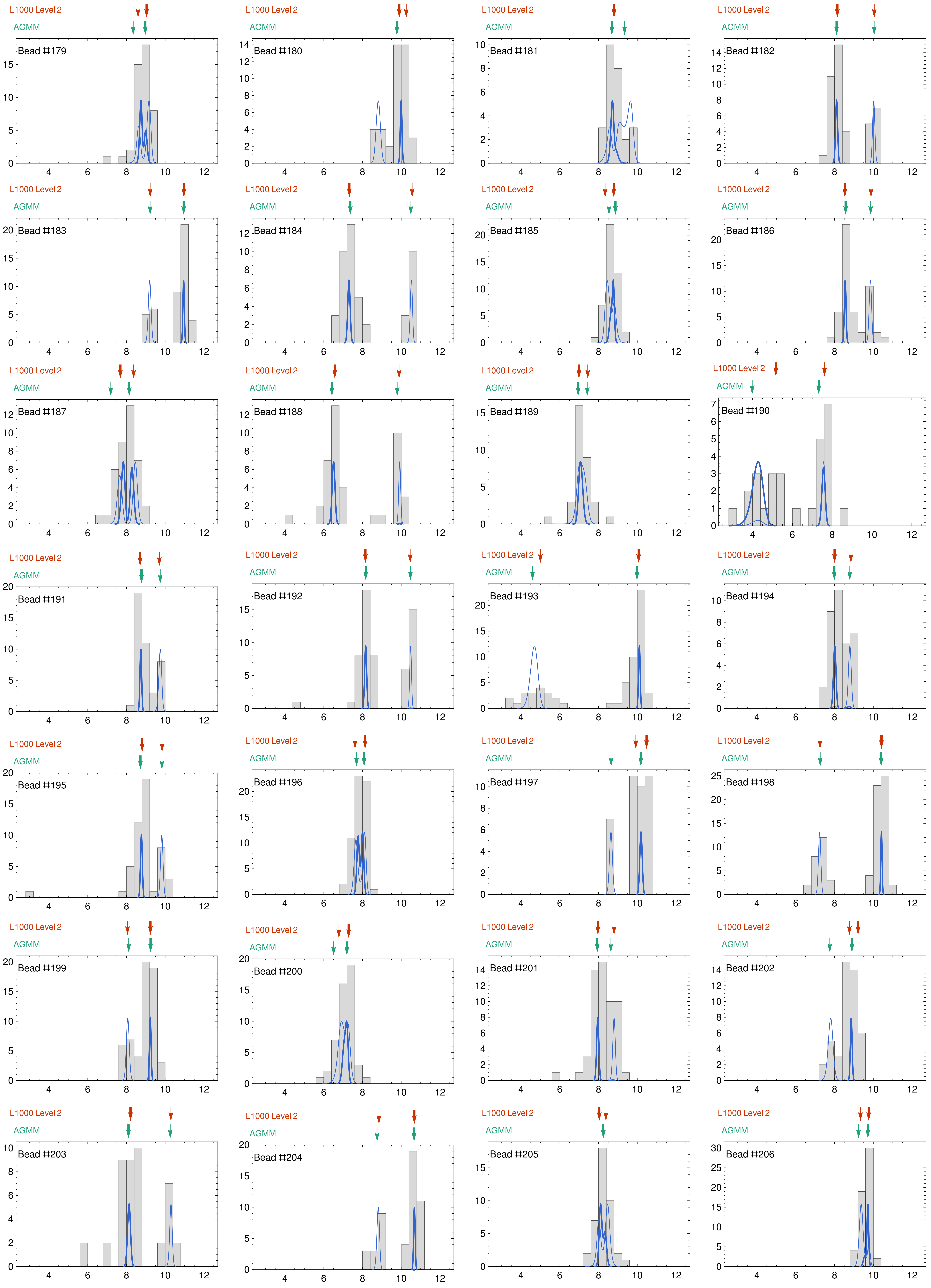
L1000 Level 2
AGMM
Bead #179
Bead #180
Bead #181
Bead #182
Bead #183
Bead #184
Bead #185
Bead #186
Bead #187
Bead #188
Bead #189
Bead #190
Bead #191
Bead #192
Bead #193
Bead #194
Bead #195
Bead #196
Bead #197
Bead #198
Bead #199
Bead #200
Bead #201
Bead #202
Bead #203
Bead #204
Bead #205
Bead #206

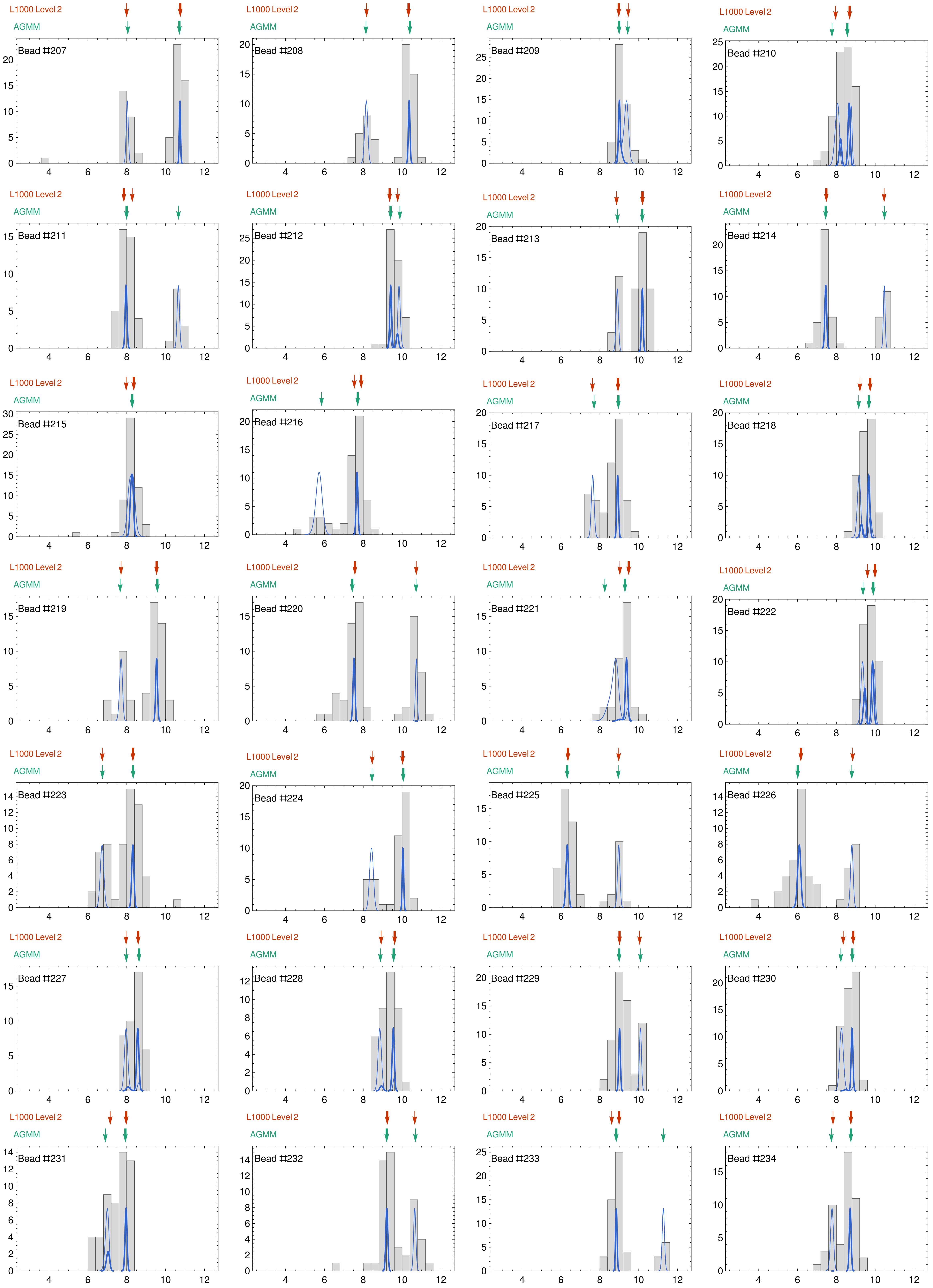
L1000 Level 2
AGMM
Bead ⌗207
Bead ⌗208
Bead ⌗209
Bead ⌗210
Bead ⌗211
Bead ⌗212
Bead ⌗213
Bead ⌗214
Bead ⌗215
Bead ⌗216
Bead ⌗217
Bead ⌗218
Bead ⌗219
Bead ⌗220
Bead ⌗221
Bead ⌗222
Bead ⌗223
Bead ⌗224
Bead ⌗225
Bead ⌗226
Bead ⌗227
Bead ⌗228
Bead ⌗229
Bead ⌗230
Bead ⌗231
Bead ⌗232
Bead ⌗233
Bead ⌗234

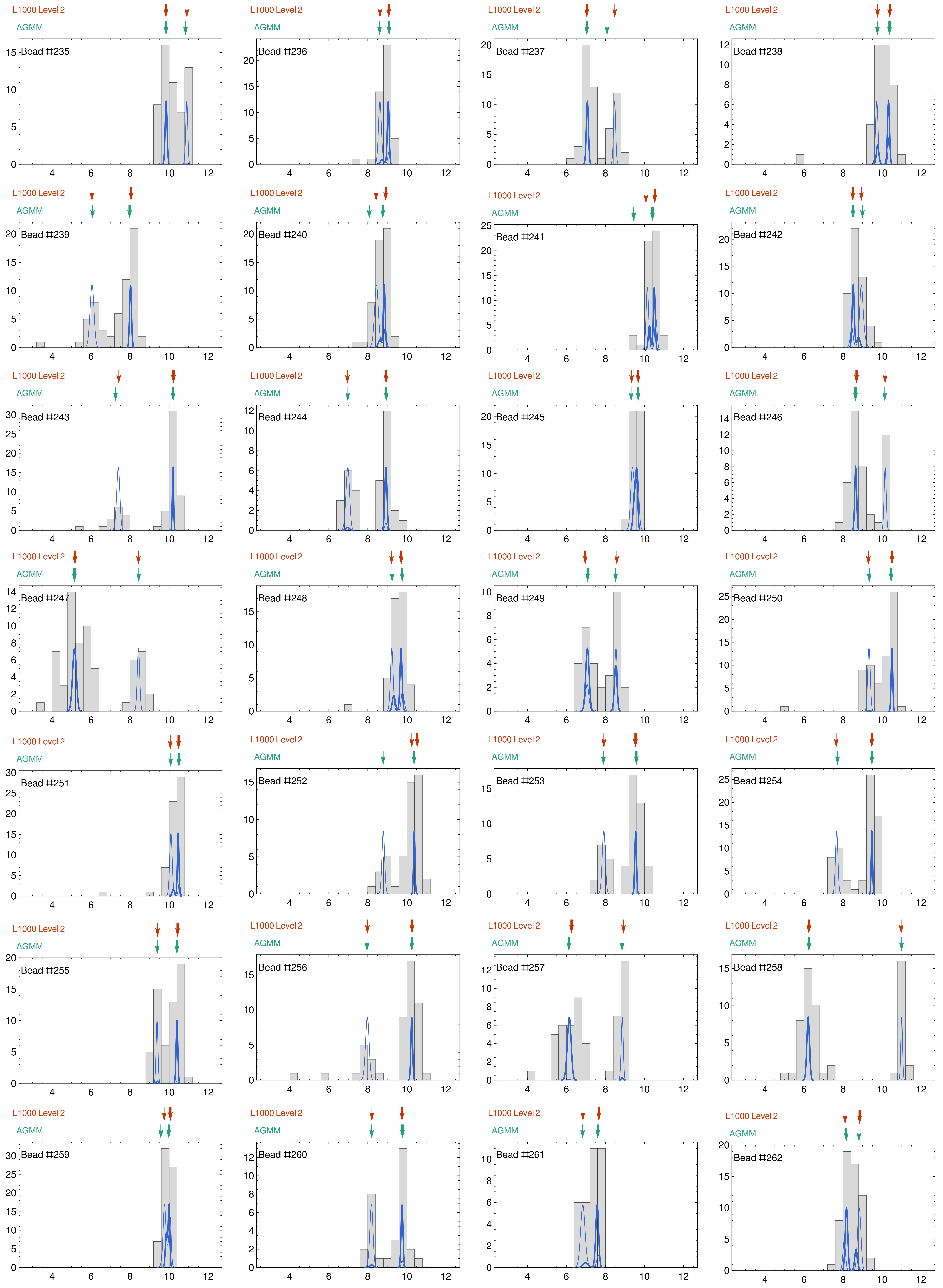
L1000 Level 2
AGMM
Bead #235
Bead #236
Bead #237
Bead #238
Bead #239
Bead #240
Bead #241
Bead #242
Bead #243
Bead #244
Bead #245
Bead #246
Bead #247
Bead #248
Bead #249
Bead #250
Bead #251
Bead #252
Bead #253
Bead #254
Bead #255
Bead #256
Bead #257
Bead #258
Bead #259
Bead #260
Bead #261
Bead #262

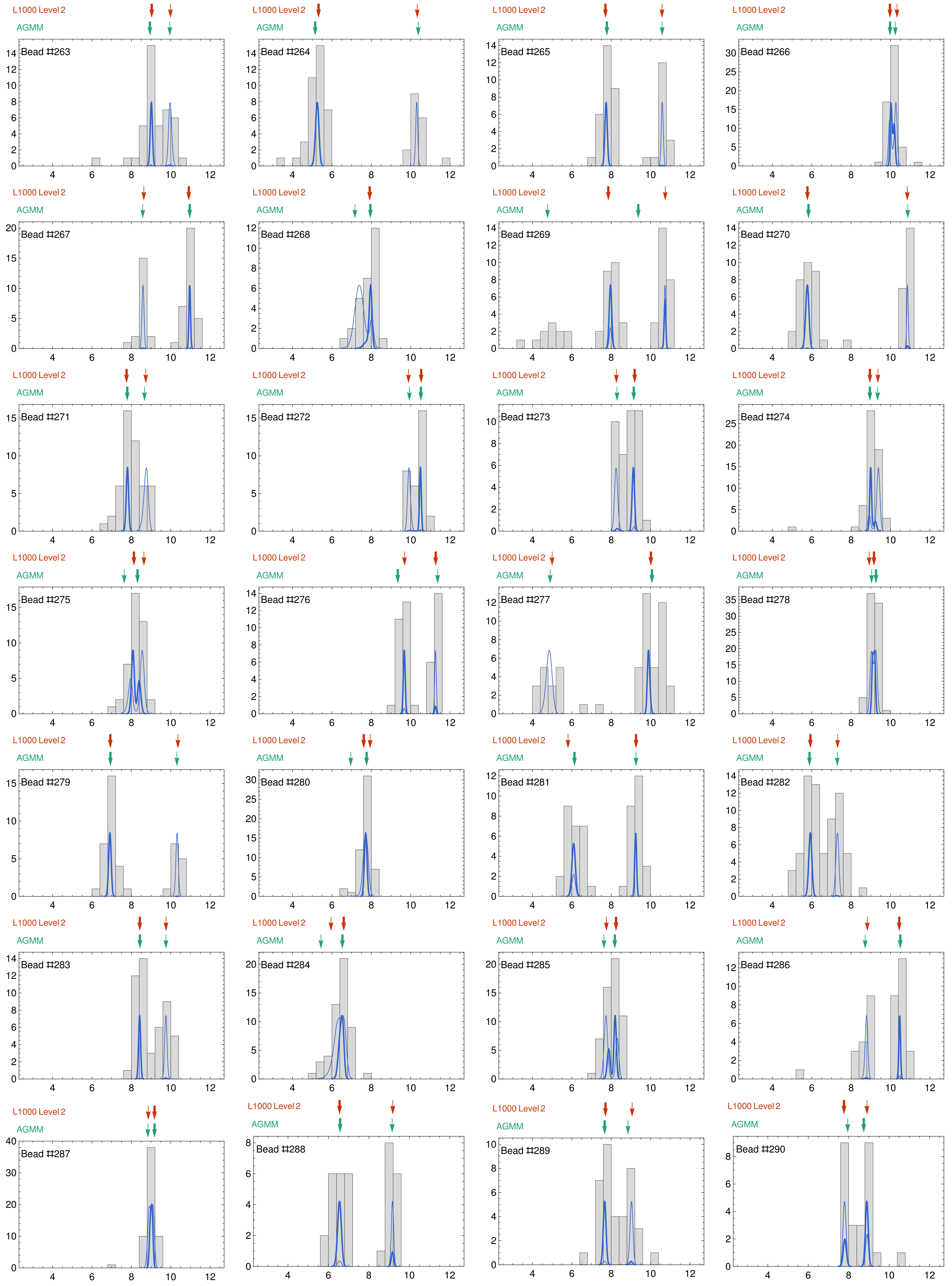
L1000 Level 2
AGMM
Bead #263
Bead #264
Bead #265
Bead #266
Bead #267
Bead #268
Bead #269
Bead #270
Bead #271
Bead #272
Bead #273
Bead #274
Bead #275
Bead #276
Bead #277
Bead #278
Bead #279
Bead #280
Bead #281
Bead #282
Bead #283
Bead #284
Bead #285
Bead #286
Bead #287
Bead #288
Bead #289
Bead #290

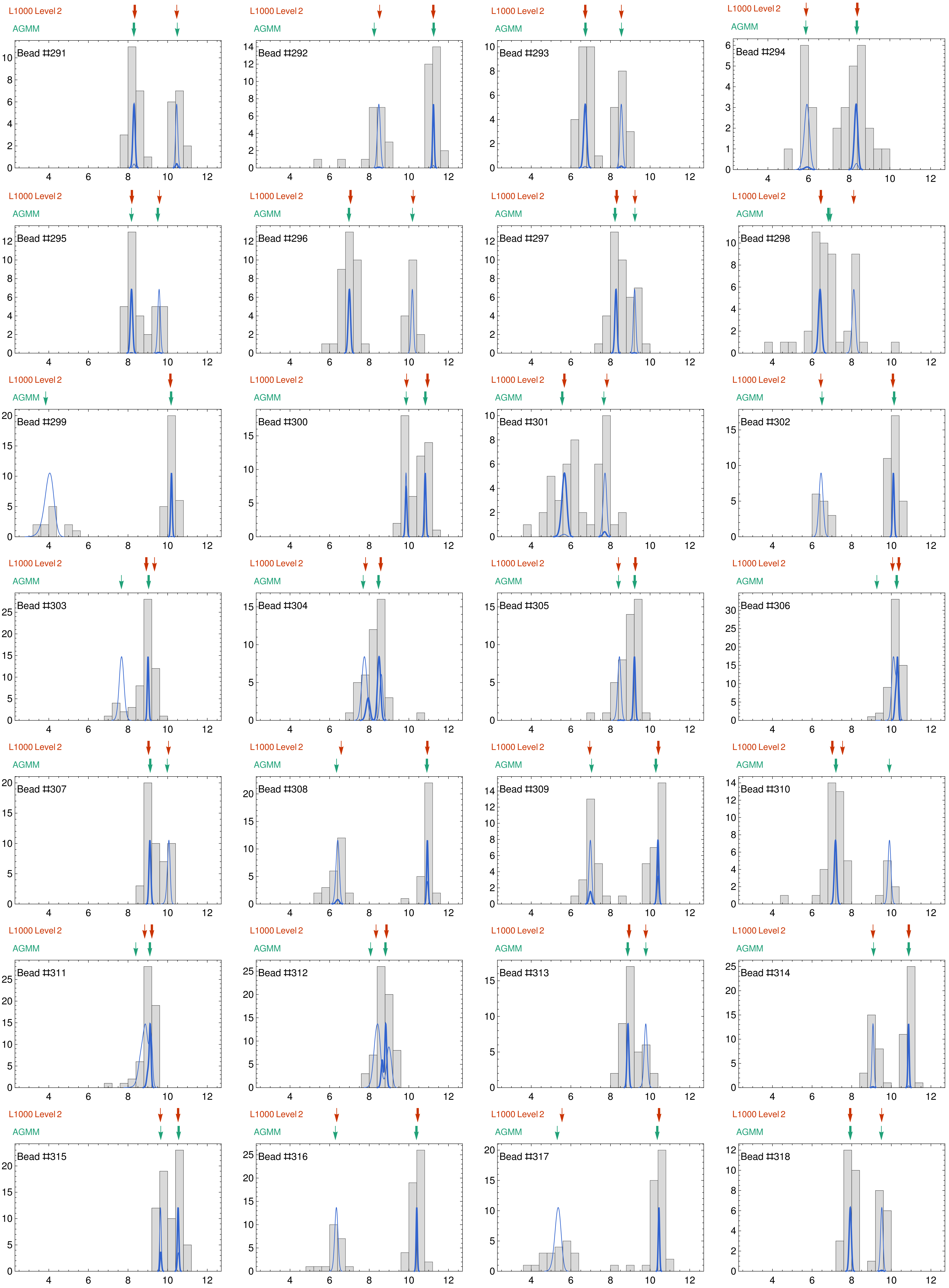
L1000 Level 2
AGMM
Bead #291
Bead #292
Bead #293
Bead #294
Bead #295
Bead #296
Bead #297
Bead #298
Bead #299
Bead #300
Bead #301
Bead #302
Bead #303
Bead #304
Bead #305
Bead #306
Bead #307
Bead #308
Bead #309
Bead #310
Bead #311
Bead #312
Bead #313
Bead #314
Bead #315
Bead #316
Bead #317
Bead #318

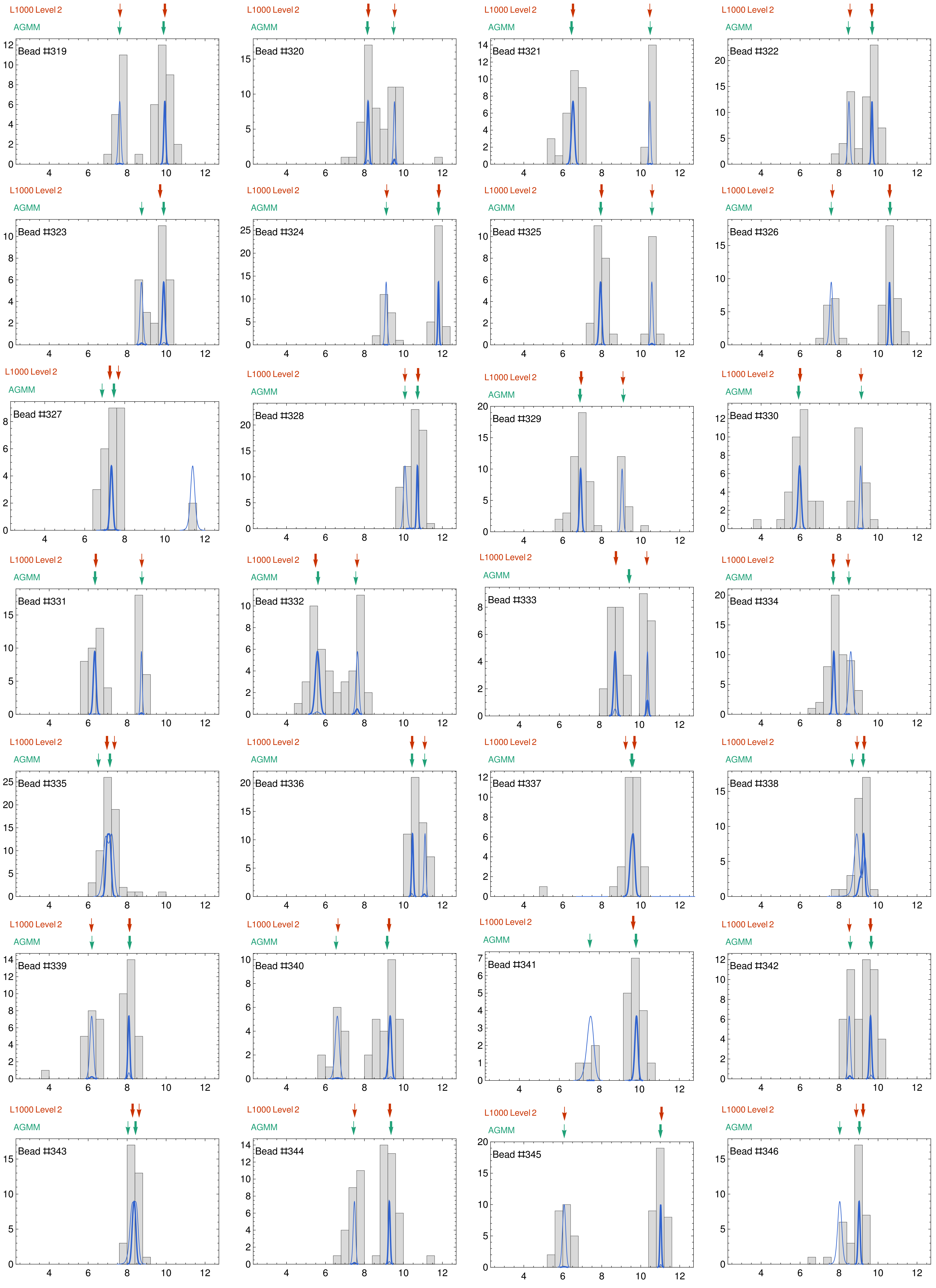
L1000 Level 2
AGMM
Bead #319
Bead #320
Bead #321
Bead #322
Bead #323
Bead #324
Bead #325
Bead #326
Bead #327
Bead #328
Bead #329
Bead #330
Bead #331
Bead #332
Bead #333
Bead #334
Bead #335
Bead #336
Bead #337
Bead #338
Bead #339
Bead #340
Bead #341
Bead #342
Bead #343
Bead #344
Bead #345
Bead #346

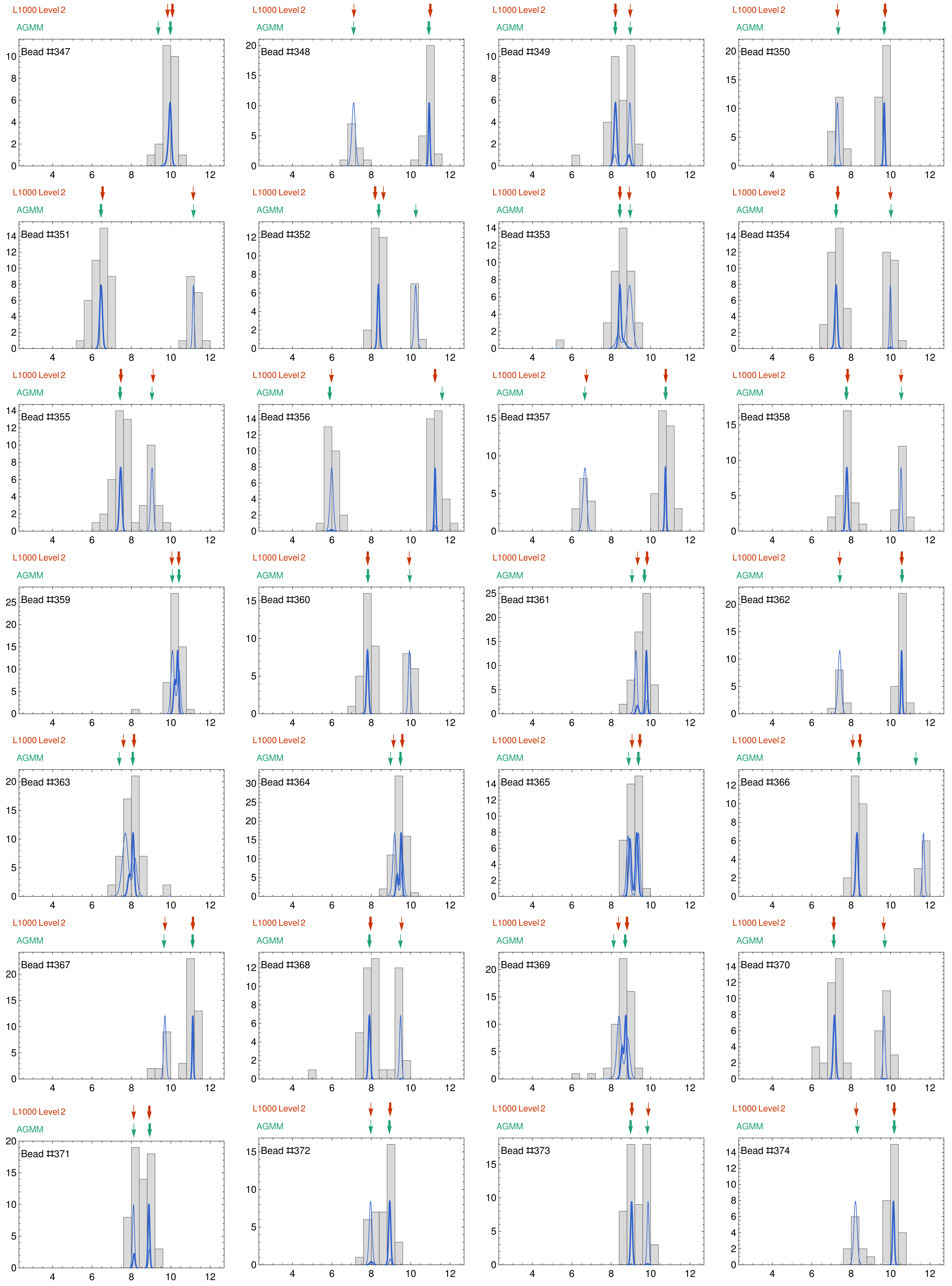
L1000 Level 2
AGMM
Bead #347
Bead #348
Bead #349
Bead #350
Bead #351
Bead #352
Bead #353
Bead #354
Bead #355
Bead #356
Bead #357
Bead #358
Bead #359
Bead #360
Bead #361
Bead #362
Bead #363
Bead #364
Bead #365
Bead #366
Bead #367
Bead #368
Bead #369
Bead #370
Bead #371
Bead #372
Bead #373
Bead #374

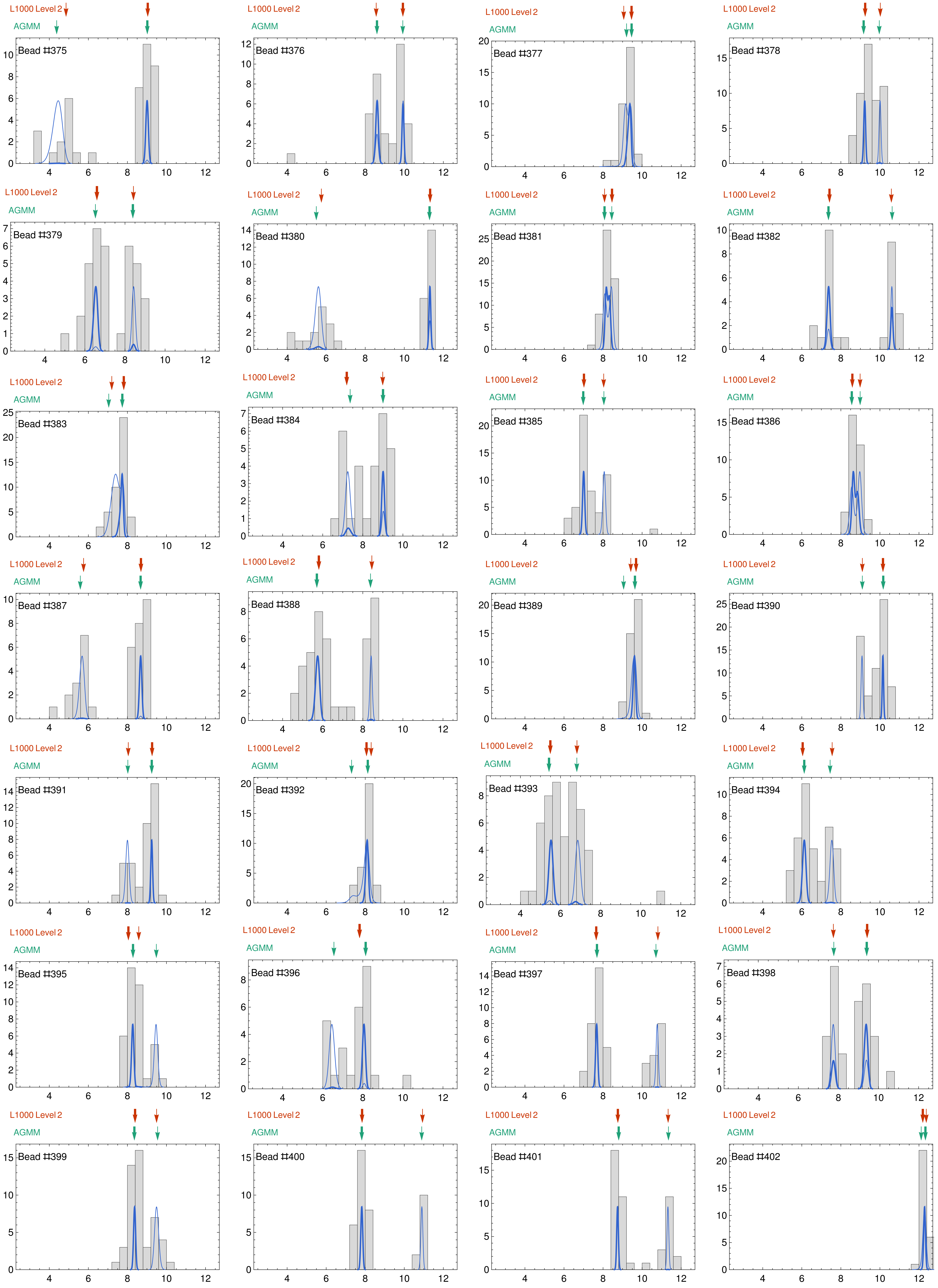
L1000 Level 2
AGMM
Bead #375
L1000 Level 2
AGMM
Bead #376
L1000 Level 2
AGMM
Bead #377
L1000 Level 2
AGMM
Bead #378
L1000 Level 2
AGMM
Bead #379
L1000 Level 2
AGMM
Bead #380
L1000 Level 2
AGMM
Bead #381
L1000 Level 2
AGMM
Bead #382
L1000 Level 2
AGMM
Bead #383
L1000 Level 2
AGMM
Bead #384
L1000 Level 2
AGMM
Bead #385
L1000 Level 2
AGMM
Bead #386
L1000 Level 2
AGMM
Bead #387
L1000 Level 2
AGMM
Bead #388
L1000 Level 2
AGMM
Bead #389
L1000 Level 2
AGMM
Bead #390
L1000 Level 2
AGMM
Bead #391
L1000 Level 2
AGMM
Bead #392
L1000 Level 2
AGMM
Bead #393
L1000 Level 2
AGMM
Bead #394
L1000 Level 2
AGMM
Bead #395
L1000 Level 2
AGMM
Bead #396
L1000 Level 2
AGMM
Bead #397
L1000 Level 2
AGMM
Bead #398
L1000 Level 2
AGMM
Bead #399
L1000 Level 2
AGMM
Bead #400
L1000 Level 2
AGMM
Bead #401
L1000 Level 2
AGMM
Bead #402

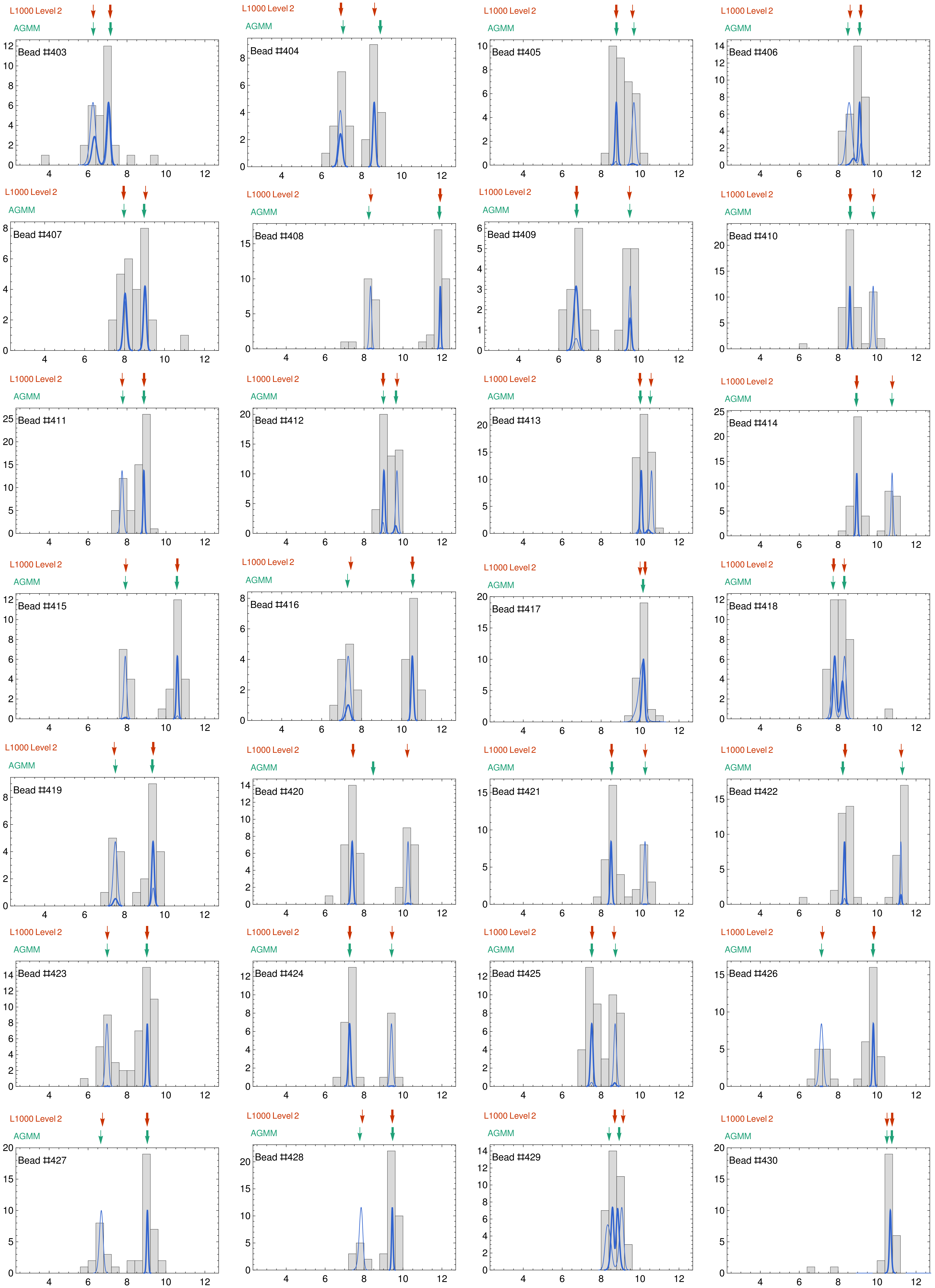
L1000 Level 2
AGMM
Bead #403
Bead #404
Bead #405
Bead #406
Bead #407
Bead #408
Bead #409
Bead #410
Bead #411
Bead #412
Bead #413
Bead #414
Bead #415
Bead #416
Bead #417
Bead #418
Bead #419
Bead #420
Bead #421
Bead #422
Bead #423
Bead #424
Bead #425
Bead #426
Bead #427
Bead #428
Bead #429
Bead #430

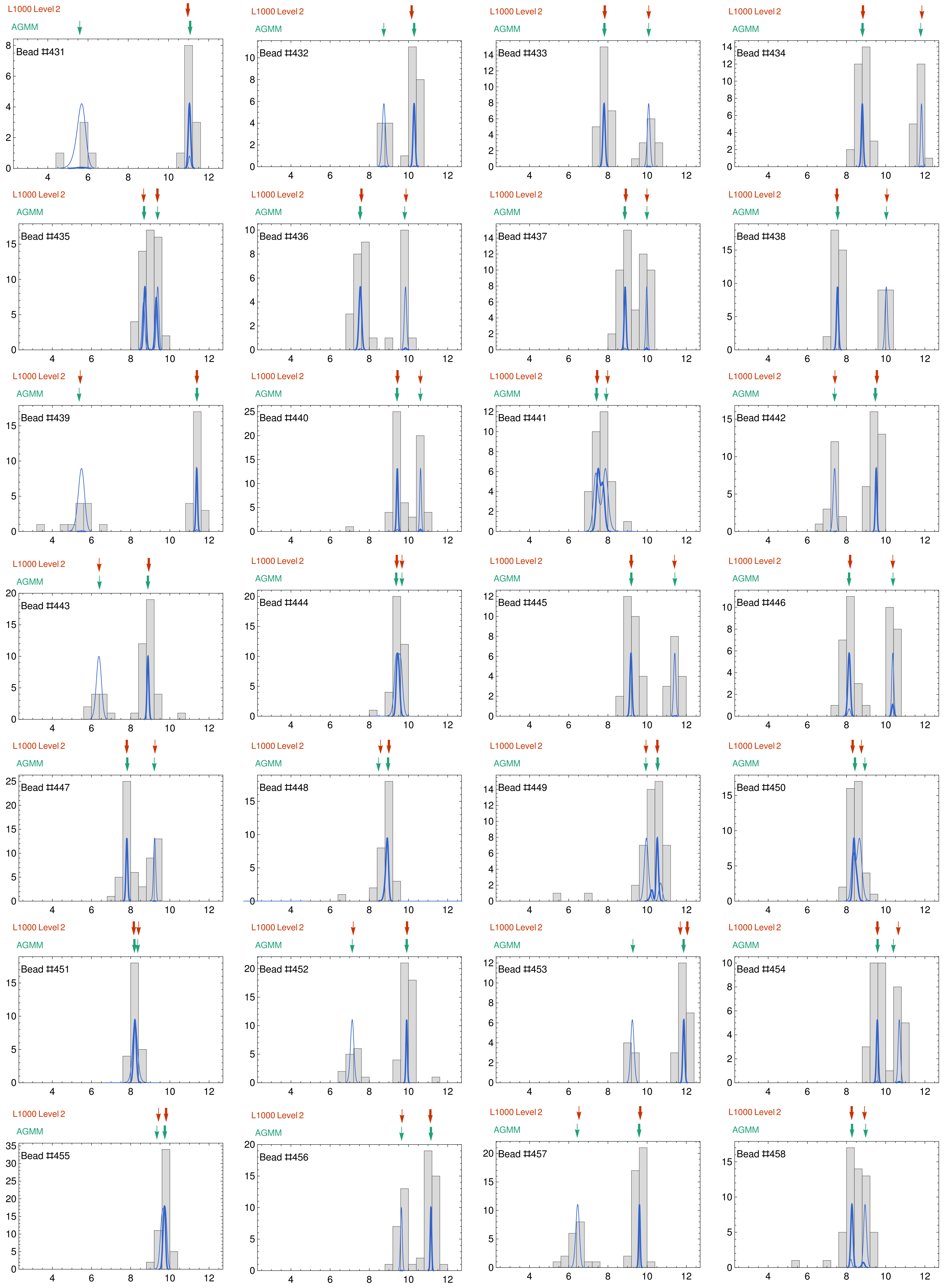
L1000 Level 2
AGMM
Bead #431
Bead #432
Bead #433
Bead #434
Bead #435
Bead #436
Bead #437
Bead #438
Bead #439
Bead #440
Bead #441
Bead #442
Bead #443
Bead #444
Bead #445
Bead #446
Bead #447
Bead #448
Bead #449
Bead #450
Bead #451
Bead #452
Bead #453
Bead #454
Bead #455
Bead #456
Bead #457
Bead #458

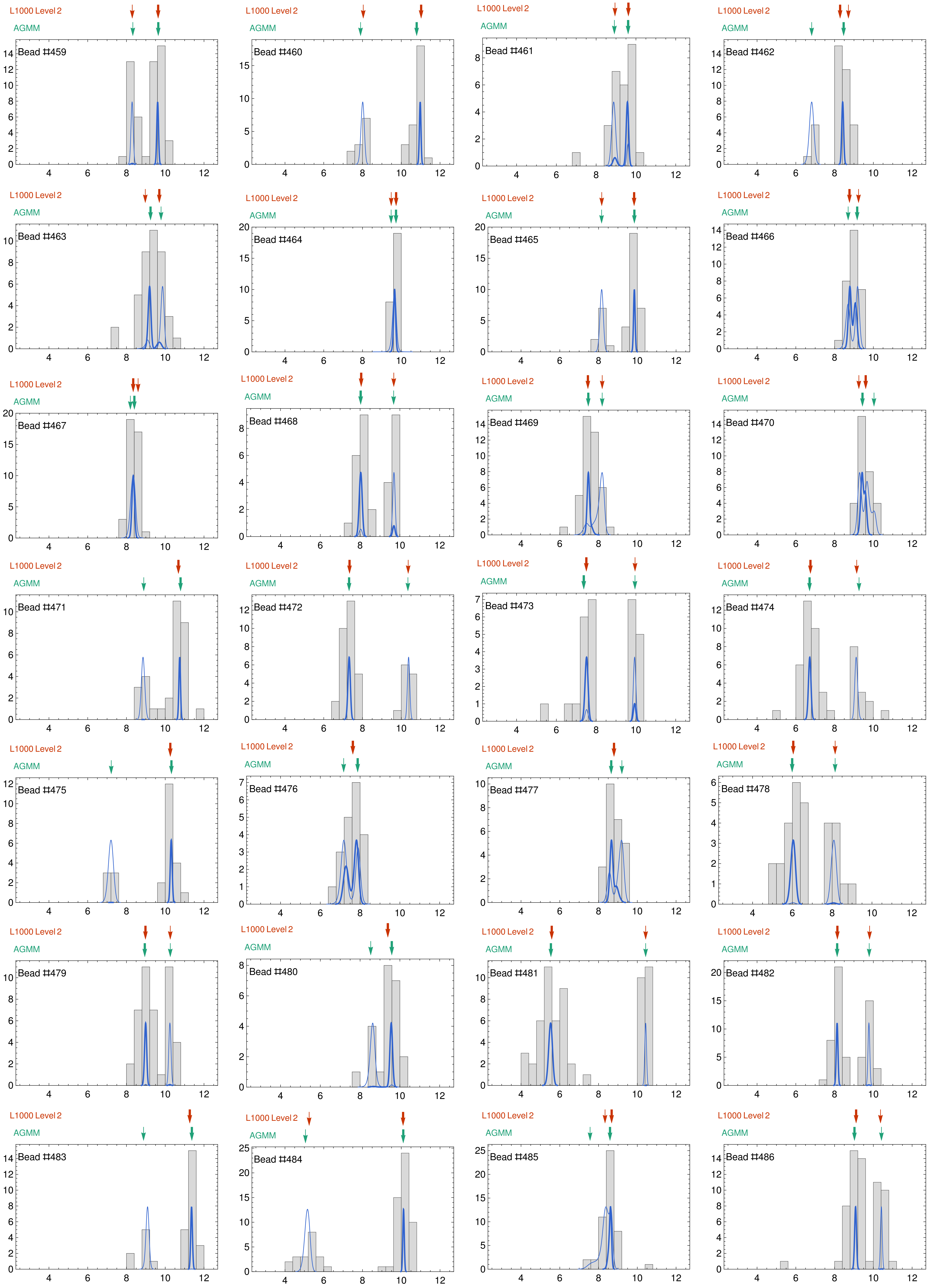

L1000 Level 2
AGMM
Bead #459
Bead #460
Bead #461
Bead #462
Bead #463
Bead #464
Bead #465
Bead #466
Bead #467
Bead #468
Bead #469
Bead #470
Bead #471
Bead #472
Bead #473
Bead #474
Bead #475
Bead #476
Bead #477
Bead #478
Bead #479
Bead #480
Bead #481
Bead #482
Bead #483
Bead #484
Bead #485
Bead #486

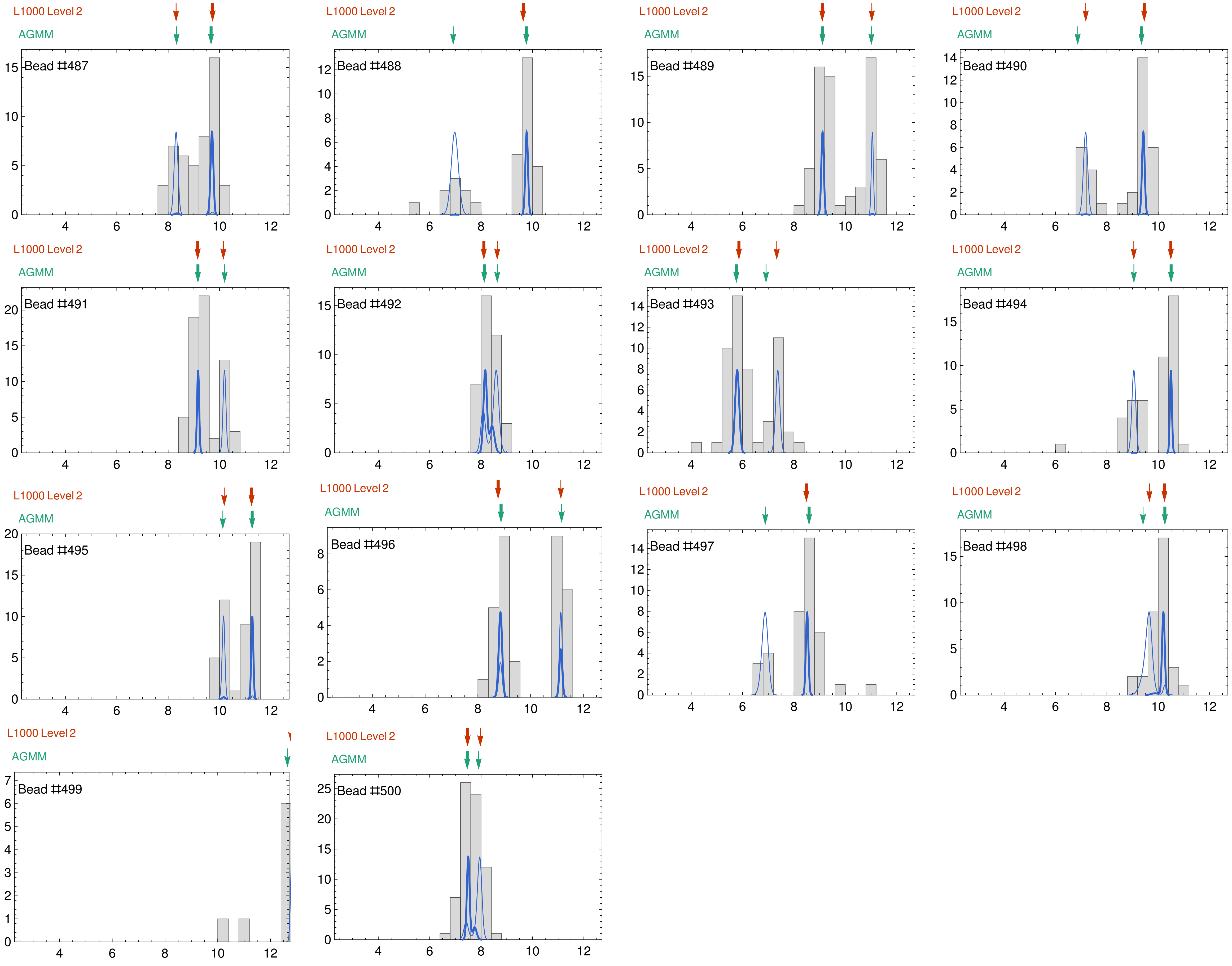

L1000 Level 2
AGMM
Bead ♯487
Bead ♯488
Bead ♯489
Bead ♯490
Bead ♯491
Bead ♯492
Bead ♯493
Bead ♯494
Bead ♯495
Bead ♯496
Bead ♯497
Bead ♯498
Bead ♯499
Bead ♯500
